# Investigating The Role Of Molecular Coating in Human Corneal Endothelial Cell Primary Culture using Artificial Intelligence-driven image analysis

**DOI:** 10.1101/2025.04.15.648927

**Authors:** Gauthier Travers, Louise Coulomb, Inès Aouimeur, Zhiguo He, Guillaume Bonnet, Edouard Ollier, Yann Gavet, Anaick Moisan, Philippe Gain, Gilles Thuret, Corantin Maurin

**Affiliations:** Laboratory Biology, Engineering and Imaging for Ophthalmology (BiiO), Faculty of Medicine, Health Innovation Campus, Jean Monnet University, Saint-Étienne, France; Institut Mines Telecom, Georges Friedel Laboratory (LGF) UMR CNRS 5307, Saint-Étienne, France; Clinical Research Unit, University Hospital, Saint-Étienne, France; Cell Therapy Unit, Auvergne Rhône Alpes French Blood Center, Saint-Ismier, France; Ophthalmology Department, University Hospital, Saint-Étienne, France

**Author notes:** Correspondence: Corantin MAURIN, PhD, Laboratory Biology, engineering and imaging for Ophthalmology (BiiO), Faculty of Medicine, Health Innovation Campus, Jean Monnet University, 10 rue de la Marandière, 42270 Saint-Priest en Jarez, France. equivalent author (Gauthier Travers, Louise Coulomb).

**Keywords:** tissue-engineered endothelial keratoplasty (TEEK), cell therapy, human corneal endothelial cells (hCECs), primary cell culture, immunohistochemistry, endothelial cell density (ECD), molecular coating, high-performance automatic segmentation

## Abstract

The monolayer of approximately 300,000 human corneal endothelial cells (hCECs) on the posterior surface of the cornea is essential to maintain transparency but is non-self-regenerative. Corneal blindness can currently only be treated by corneal transplantation, hindered by a global donor shortage, highlighting the need for developing tissue and/or cell therapy. The mass production of these advanced therapy medicinal products requires obtaining high-yield, high-quality endothelial cell culture characterized by hexagonal shape, low size variability, and high endothelial cell density (ECD). Among the usual critical quality attributes which combine the expression of differentiation markers, ECD and cell morphology parameters, the latter are not optimally measured *in vitro* by conventional image analysis which poorly recognize adherent cultured cells. We developed a high-performance automated segmentation using Cellpose algorithm and an original analysis method, improving calculation of classical morphological parameters (coefficient of variation of cell area and hexagonality) and introducing new parameters specific to hCECs culture *in vitro*. Considering the importance of the extracellular matrix *in vivo*, and the panel of molecules available for coating cell culture plastics, we used these new tools to perform a comprehensive comparison of 13 molecules (laminins and collagens). We demonstrated their ability to discriminate subtle differences between cultures.

## 1. Introduction

Human Corneal Endothelial Cells (hCECs) form a monolayer of hexagonal cells at the inner part of the cornea, resting on the Descemet’s membrane (DM). They have a crucial role of barrier and ion pump function to ensure deswelling of the hydrophilic stroma. This function is essential for maintaining corneal thickness and transparency. Corneal endothelial dysfunction is the major cause of corneal low vision worldwide [1], [2]. The only treatment for these diseases is corneal transplantation with around 185000 keratoplasty per year worldwide. However, the global abysmal chronic shortage of donors highlights the importance of developing alternative solutions [3]. Cell therapy is now a reality in Japan, where it has been approved by health authorities, and may soon be in the United States of America. Complementary approaches, such as tissue engineering, could also be useful to meet this growing demand [4], [5], [6]. For both advanced therapy medicinal products (ATMP), the production of high-quality clinical grade hCECs *in vitro* (small, hexagonal and cohesive) is essential and particularly challenging because of the low regenerative capacity already mentioned [7]. So far, all hCECs cultures of clinical-grade have been exclusively performed using donors under the age of 35, as their cells are more numerous and exhibit significantly higher proliferative capacities compared to older donors [8], [9], [10]. However, these young donors are extremely rare due to a very high refusal rate for pediatric donations, and the (fortunately) low mortality among young adults.

A persistent challenge is therefore to establish high yield culture of hCECs sourced from older donors using innovative methods to increase hCECs density *in vitro* while avoiding pheno-typic changes such as endothelial mesenchymal transition (EndoMT) [11], [12], [13], [14]. In EndoMT, cells lose their hexagonal-shape and intercellular junction, increase their production of extracellular matrix (ECM), change their gene expression and, hypothetically, their functionality (such as pump function), although this aspect is rarely investigated in most research studies. To obtain an optimal cell culture, two parameters can be improved: the culture medium and the molecular coating of culture plastics which enhances cell adherence and differentiation. The present study is focused on molecular coating.

The three main coating proteins used in hCECs culture are collagens, fibronectin and laminins [15], [16]. In Japan, the Laminin-511 E8 protein fragment has been selected as an ideal xeno-free defined substrate to produce clinical grade hCECs. For evaluating hCECs quality in adherent culture conditions, morphometric analysis has multiple advantages. Monolayer arrangement facilitates imaging, allowing for clearer observation and analysis of cellular characteristics. These culture conditions closely mimic the native tissue environment, providing a better simulation of the physiological conditions of hCECs. Therefore, *in vitro* cells morphology might be a good predictor of what will be observed in the recipient of ATMPs. In addition, it will be likely challenging to gain approval from health authorities for clinical batch release criteria if the cultured cells do not sufficiently resemble native corneal endothelium, as morphological conformity is to be a key indicator of the viability and effectiveness of cells prepared for transplantation.

Understanding cell morphology in adherent culture conditions, particularly in response to different coating molecules, could provide new insights for optimizing cell selection and improving transplantation outcomes. We hypothesize that in primary culture, a characterization based on cell morphology could potentially be used to select high-quality cells, particularly when using donors over 30 years old, who are significantly more frequent but whose hCECs are fewer in number, less proliferative and more prone to EndoMT.

In the present study, we performed a screening of several laminins and collagens to optimize hCECs quality and maximize the culture yield. Morphometry and ECD were quantified using an artificial intelligence (AI) driven segmentation tools specifically designed, trained and tuned for hCECs analysis and first validated on cultures with defined phenotypes.

## 2. Materials and Methods

### 2.1 Donor tissue and ethical statement

hCECs used in this study were obtained from primary cell cultures derived from organ-cultured human corneas. They were sourced from corneal donations for scientific purposes (laboratory of anatomy of the Faculty of Medicine of Saint-Étienne) or from three french eye banks (Besançon, Saint-Étienne, and Nantes). All corneas were unsuitable for transplantation because their parameters did not meet the required standards for clinical use. The BiiO laboratory, authorized by the Ministry of Higher Education, Research, and Innovation of France (Ministère de l’Enseignement Supérieur, de la Recherche et de l’Innovation, MESRI) under the number DC-2023-5458, conducted this research on these human corneas without the need for additional ethical approval. They were handled in accordance with the principles of the Declaration of Helsinki, bioethics laws, and French and European regulations regarding tissue donations.

For AI fine tuning, we used 15 images of 3 cultures. For AI segmentation and criteria validation, we used 4 other different endothelial cell cultures (listed in Table 1) which had varying quality, ranging from a culture undergoing endothelial-mesenchymal transition to an endothelial culture from a very young donor (2-year-old, characterized by high ECD and excellent morphology). For the comparison of coating molecules, eight cultures from eight donors were used. The donors ages ranged from 55 to 85 years, with a mean±standard deviation of 71 ± 11 years (sex ratio = 1 (4 males, 4 females) and corneas were stored at 31°C in organ culture medium (CorneaMax, Eurobio, Les Ulis, France) during 25 ± 22 days (min = 4; max = 72) before use. The initial ECD, measured within 5 days after retrieval, was 2390 ± 559 (1287– 3233) cells/mm^2^. Donors and cultures characteristics were detailed Table 1.

**Table 1.** Characteristics of donor corneas used for cell cultures.

| ID | Death-to-retrieval time (hours) | Initial ECD (cells/mm<sup>2</sup>) | Organ culture duration before primary culture (days) | Sex | Age (years) | Cell culture passage (n) | Application |
| --- | --- | --- | --- | --- | --- | --- | --- |
| $\alpha$ | 12 | 2863 | 86 | M | 58 | 9 | AI tuning |
| $\beta$ | 15 | 3248 | 6 | F | 79 | 8 | |
| $\Omega$ | 20 | 2285 | 4 | F | 59 | 6 | |
| 1 | 16 | nd | 16 | F | 91 | 7 | AI validation |
| 2 | 19 | 2395 | 50 | M | 77 | 5 |  |
| 3 | 24 | nd | 65 | M | 2 | 4 |  |
| 4 | 10 | 2874 | 26 | M | 82 | 11 |  |
| a | 23 | 2562 | 11 | F | 85 | 6 | Comparison of coating molecules |
| b | 20 | 2285 | 4 | F | 59 | 6 |  |
| c | 18 | 2154 | 13 | M | 55 | 5 |  |
| d | 1 | 1287 | 36 | F | 67 | 4 |  |
| e | 24 | 2779 | 27 | M | 79 | 4 |  |
| f | 12 | 2321 | 5 | M | 82 | 9 |  |
| g | 9 | 2495 | 29 | F | 73 | 7 |  |
| h | 11 | 3233 | 72 | M | 69 | 8 |  |

ECD = Endothelial Cell Density; nd = not determined; F = female; M = male; AI = artificial intelligence

### 2.2. Cell culture media

Three different media were used: 1/The DM digestion medium consisted of Opti-MEM (11058021, Gibco, Grand Island, NY, USA) supplemented with 200 μg/mL CaCl_2_ (C5670, Sigma-Aldrich, St. Louis, MO, USA); 20 μg/mL ascorbic acid (A5960, Sigma-Aldrich) and 2 mg/mL collagenase A (10103586001, Roche, Basel, Switzerland). 2/The basic medium consisted of Opti-MEM supplemented with antibiotic–antimycotic (152-062, Gibco) at 1/200, 8% of fetal bovin serum (S-1860-500, Eurobio), 0.1% of CaCl_2_, 0.08% of chondroitine sulfate (034-1462, FUJIFILM Wako Pure Chemical Corporation, San Diego, CA, USA), 10 μM of SB203580 (3.8 μg/mL) (S1076, SelleckChem, Houston, TX, USA) and 1 μM of SB431542 (S1067, SelleckChem) (where SB203580 was an inhibitor of p38 MAPK pathway and SB431542 an inhibitor of TGF-β receptor; both involved in the limitation of EndoMT in cell culture), 20 μg/mL of ascorbic acid, 5ng/mL of Epidermal growth factor (EGF) (PHG0311, Gibco, Grand Island, NY, USA). 3/The medium for the initiation of primary cell culture and for cell passaging was the basic medium with the addition of a rock inhibitor (RI), Y-27632 (S1049, SelleckChem) to promote cell adherence, proliferation and survival [17], [18], [19].

### 2.3. Primary culture of hCECs

hCECs were cultivated using a protocol derived from the standard peel and digest method, also used by the team which selected the coating with iMatrix-511 (892012, Nippi, Incorporated, Tokyo, Japan) [20]. We increased the digestion time from 12 to 16 hours and used a higher collagenase concentration (1 mg/mL to 2 mg/mL) to enhance cell dissociation as previously described [21]. To isolate hCECs, DM was mechanically peeled from donor corneas, ensuring maximum peripheral coverage. The membranes were then incubated at 37°C in a humidified 5% CO_2_ atmosphere for 16 hours with 0.5 mL of digestion medium in 48-well plates (3548, Corning, Kennebunk, ME, USA). After digestion, released hCECs were centrifuged at 200g for 5 minutes in conical-bottom tubes (641997, Dutscher, Caplugs Evergreen, CA, USA). The cells were then seeded in 24-well plates (3524, Corning) pre-coated with iMatrix-511 according to the manufacturer’s instructions. hCECs were cultured in a humidified 5% CO_2_ atmosphere at 37°C. The basic medium was changed the day after seeding and then weekly until confluence, which took about a month.

For cell passage, hCECs culture were first rinsed in Ca^2+^ and Mg^2+^-free phosphate buffered saline (PBS) (SH30256.02, Cytiva, MA, USA), then incubated in TripLE Select Enzyme 5X (A1217701, Gibco) during 15 minutes at 37°C until cell detachment. Cells were detached mechanically by aspirating and dispensing the liquid in the well and chemically using TrypLE solution. Cells were then transferred in conical-bottom tubes containing basic medium supplemented with RI. The complete detachment of cells was confirmed using a phase-contrast microscope (CKX41, Olympus, Tokyo, Japan). Viable cells were counted using Trypan blue staining and an automatic cell counter (TC10, Bio-Rad, Hercules, CA, USA). Counting was performed twice to ensure repeatability and to calculate the average cell number. Passage followed a defined scaling ratio: P1 (1 well of a 24-well plate into 1 well of a 12-well plate), and P2 (1 well of a 12-well plate into 1 well of a 6-well plate), with subsequent passages at a 1:2 ratio. Throughout all stages, cells used for experiments were cultured on plates coated with the reference molecule, iMatrix-511 until experiment with the different molecular coatings. Cultured hCECs at passages 4 through 9 were used for these experiments. After the last cell counting, hCECs were centrifuged at 200g for 5 minutes. The supernatant was removed and hCECs were resuspended in the medium previously described and seeded, this time, at 500 cells/mm^2^ in 384 wells plates (353961, Falcon, MA, USA) for experiments with the different molecular coatings.

To validate our AI image analysis method, we selected four cultures with very different phenotypes, based on conventional naked-eye microscopy observations, representative of the various endothelial qualities, ranging from the poorest (non-transplantable) to the best (the desired goal, i.e. deemed suitable for clinical application), as illustrated in Fig. 1. Culture 1 underwent EndoMT. Culture 2 was typical of a double population of normal and likely senescent cells. Culture 3 and 4 both were clinically acceptable compared to the initial clinical trial [9]. For this step, each of the 4 cultures was cultured on plates coated with iMatrix-511 (the referenced coating) in 384 wells with 16 replicates per culture.

**Fig. 1.**
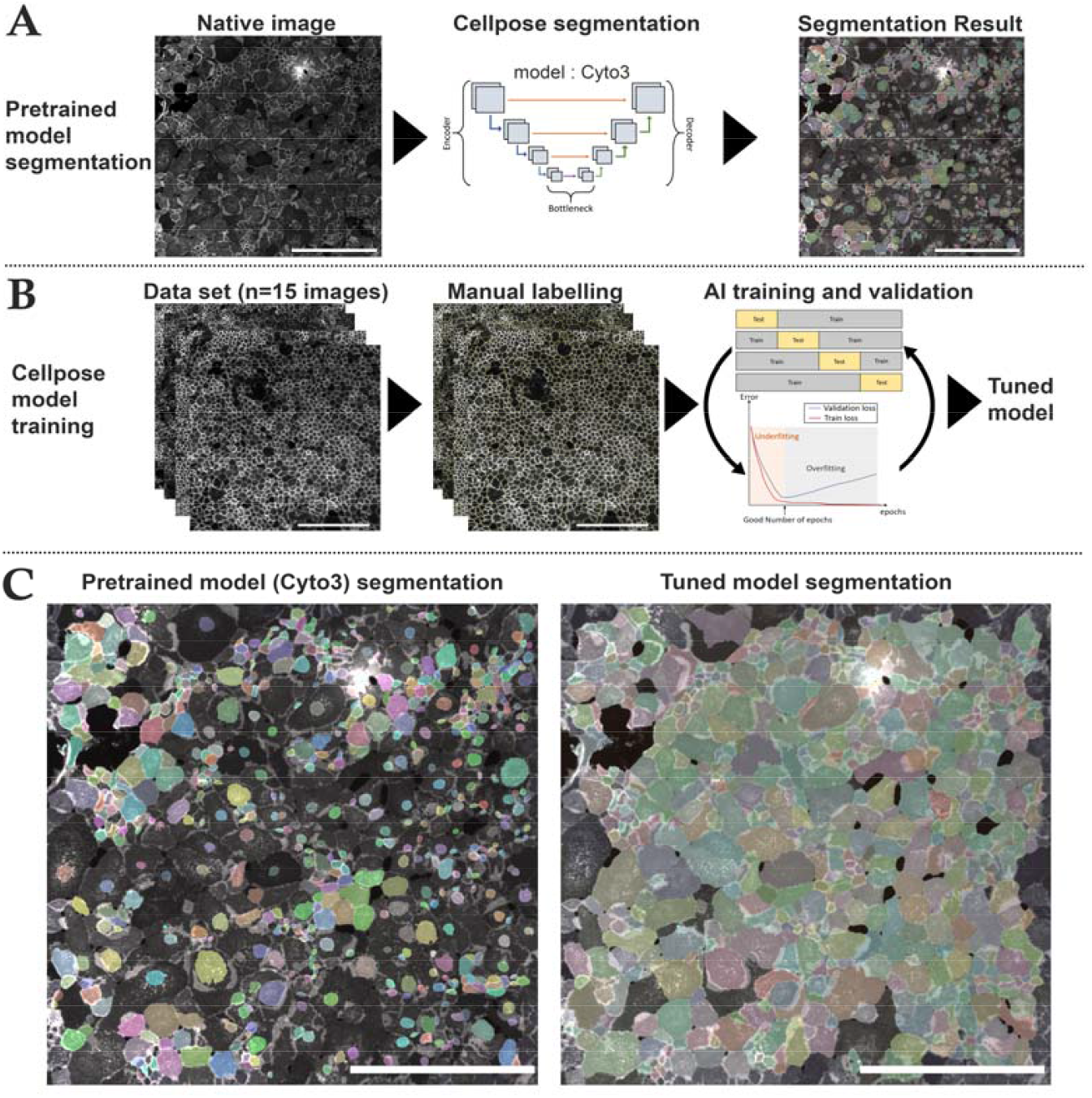
Cell segmentation and AI training steps using cellpose. **(A)** Steps for image segmentation with cellpose which used an U-net model pre-trained on different cell types (Cyto3 model). The initial segmentation was not efficient on NCAM-labelled cultured corneal endothelial cells. Cell areas were coded by colors. **(B)** Steps for training a new model consisted in the selection of NCAM-labelled image of various quality (n=15 images), followed by manual labelling and then a 4-fold cross validation training to optimize epochs and control segmentation quality of the newly trained model (tuned model). **(C)** Comparison of the same image segmented with the pre-trained Cyto3 model and with our tuned model. Scale bar = 500μm on each image.

In a second step, we used this validated AI image analysis method to evaluate 13 coating molecules on 8 different cultures. The diversity of the parameters involved, particularly the inter-donor variability and the variability associated with the coating molecules, made it impossible to subjectively discriminate by the naked eye, which justified the use of an automated analysis method. The 8 cultures were selected based on their ECD higher than 1000 cells/mm^2^. This rather low threshold (by reference to the usual eye bank lower threshold of 2000 cells/mm^2^ for transplantation) was chosen because the aim of this work was to screen for conditions liable to increase final ECD.

For the screening of molecular coating, for each coating condition tested, 8 replicates were performed for each of the 8 different corneal donors (1 coating molecule, 1 donor, 8 repetitions). We selected 9 laminin isoforms (all from BioLamina, Sundbyberg, Sweden) as well as iMatrix-511 (Nippi) and 3 types of collagens (all from Sigma Aldrich) (Table 2). Laminins differed in their chain composition, tissue localization, and biological functions, making them of interest for specific applications in regenerative medicine and tissue engineering. These 13 molecules were compared to a negative control (uncoated well). The coating with the ten laminin molecules, including the control, was performed overnight at 4°C, as recommended by the supplier, with all at 1 μg/cm^2^ except for iMatrix-511, which was coated at 0.5 μg/cm^2^. For the three collagen molecules, incubation time was done at 37°C until complete drying of the well. Collagens I and IV were performed at 6 μg/cm^2^ and collagen II at 2 μg/cm^2^.

**Table 2.**
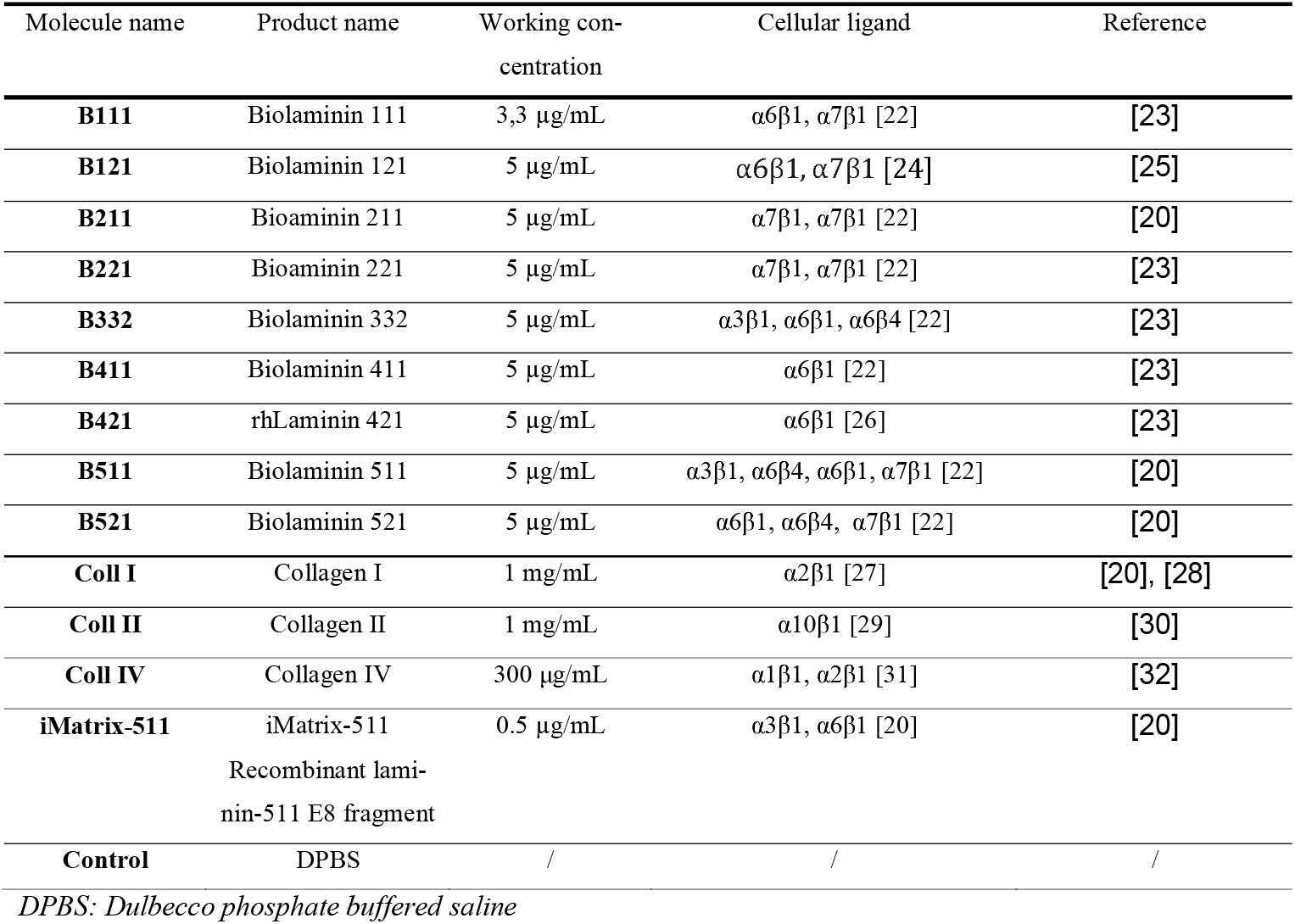
Characteristics of the nine molecular coating.

### 2.5. Immunocytochemistry

At confluency, after approximately 3 weeks of culture, immunocytochemistry (ICC) was performed on hCECs directly in culture plates. The ICC protocol was previously described [21], [33]. Briefly, cells were rinsed in Ca^2+^ and Mg^2+^ DPBS (SH30028.FS, Cytiva, Saint-Germain-en-Laye, France) and then fixed in pure methanol for 15 minutes at room temperature (RT). After three rinses, with Ca^2+^ and Mg^2+^ DPBS, cells were incubated in saturation buffer composed of 2% bovine serum albumin (A3059-100G, Sigma-Aldrich) and 2% goat serum (191356, MP Biomedicals, Irvine, CA, USA) for 30 minutes at 37°C. The hCECs were incubated with primary antibody for 1 hour at 37°C: NCAM (1:400, MAB24081, R&D system, Minneapolis, MN, USA). After three rinses in PBS, the highly cross-absorbed secondary antibodies (Alexa Fluor 488-conjugated goat anti-rabbit IgG (A11034, Invitrogen, Carlsbad, CA, USA)), diluted to 1/800, and DAPI diluted at 2 μg/mL in blocking buffer were incubated for one hour at 37°C under gentle agitation. After rinsing three times, Fluoromount-G mounting medium (00-4958-02, Invitrogen) was added in each well.

### 2.6. Images acquisition

The samples were observed with an inverted fluorescence microscope IX81 (Olympus), equipped with CellSens Dimension software V2.3 (Olympus). Images were taken at x10 and x40 magnification on the center of each well. Images were obtained using a FITC and DAPI filter, using optimized parameters for each image (intensity of light source, exposure time, contrast, resolution) and a CMOS camera 2024x2048 pixels (Orcas-Flash4.0 LT+, C11440-42U30, Hamamatsu photonics K.K., Hamamatsu city, Japan).

### 2.7. Endothelial cell density measurement

After staining, 8 replicate images per molecular coating were taken for each donor, to mesure ECD. For this, we automated the counting of each cell nuclei (DAPI stained) with a macro using the “Stardist” plugin (https://github.com/stardist/stardist) on Fiji Software (V2.9.0/1.53t https://fiji.sc/) using x10 magnification images [34], [35]. The software segmented each stained nuclei and calculated the ECD according to the pixel ratio. We performed a visual inspection to ensure proper nuclear recognition and corrected any possible errors.

### 2.8. Cellular segmentation and morphological analysis using AI and mathematical algorithms

To characterize hCECs morphology, we developed a high-performance AI based automatic segmentation and analysis solution for cells labeled with a lateral membrane marker (here NCAM) and a nucleus marker (DAPI). For morphological analysis, 2 images were taken per molecular coating and for each donor.

#### 2.8.1. Segmentation using AI

We used Cellpose, an open source generalist deep learning algorithm for cell and nucleus segmentation (https://github.com/MouseLand/cellpose) that can be tuned for optimizing results with the users data (Fig. 1) [36]. We selected the cyto3 model because of its versatility and better efficacy than other available pretrained models [37]. We first verified if it was efficient with our images. We then tuned the “cyto3” Cellpose model with hCECs images labeled with NCAM and DAPI. We manually segmented 15 images of different cell culture quality directly on the cellpose Graphical User Interface (GUI) which allowed extracting the result of the segmentation on a separate image as well as the regions of interest (ROI) of each image. Manual segmentation took approximately 4-8 hours per image depending on image complexity and ECD.

We then performed a 4-fold cross-validation on the python version of cellpose based on the 15 manually segmented images (2048x2048 pixel for 1330x1330μm, 16 bits) coupled with the fluorescence images (membranes and nuclei in two different image channels). Data were divided into 60% training set, 20% test set and 20% validation set, data augmentation was performed with a factor of 4 by horizontally and vertically flipping images. As there were approximately 1500 cells per image, the training set was thus constituted of around 54000 cells. For each fold, we optimized epochs (number of training cycles the AI went through to learn/correct the weights of the model) using the validation set (800 was the best epochs per fold). Cross validation was done by comparing manually segmented images with Cellpose segmentation using a published dissimilarity criterion and the 6 parameters described hereafter (ECD, CV, adjusted CV, HEX, HEX-Q, Filimorphism) [38].

#### 2.8.2. Morphometric analysis

We developed a custom script in python language (V3.11.4, https://www.python.org/downloads/release/python-3114/) interpreted with Spyder IDE (V5.4.3) to measure 5 parameters on segmented images, to conduct a comprehensive analysis of cell polymegathism and pleomorphism (Fig. 2):

**Fig. 2.**
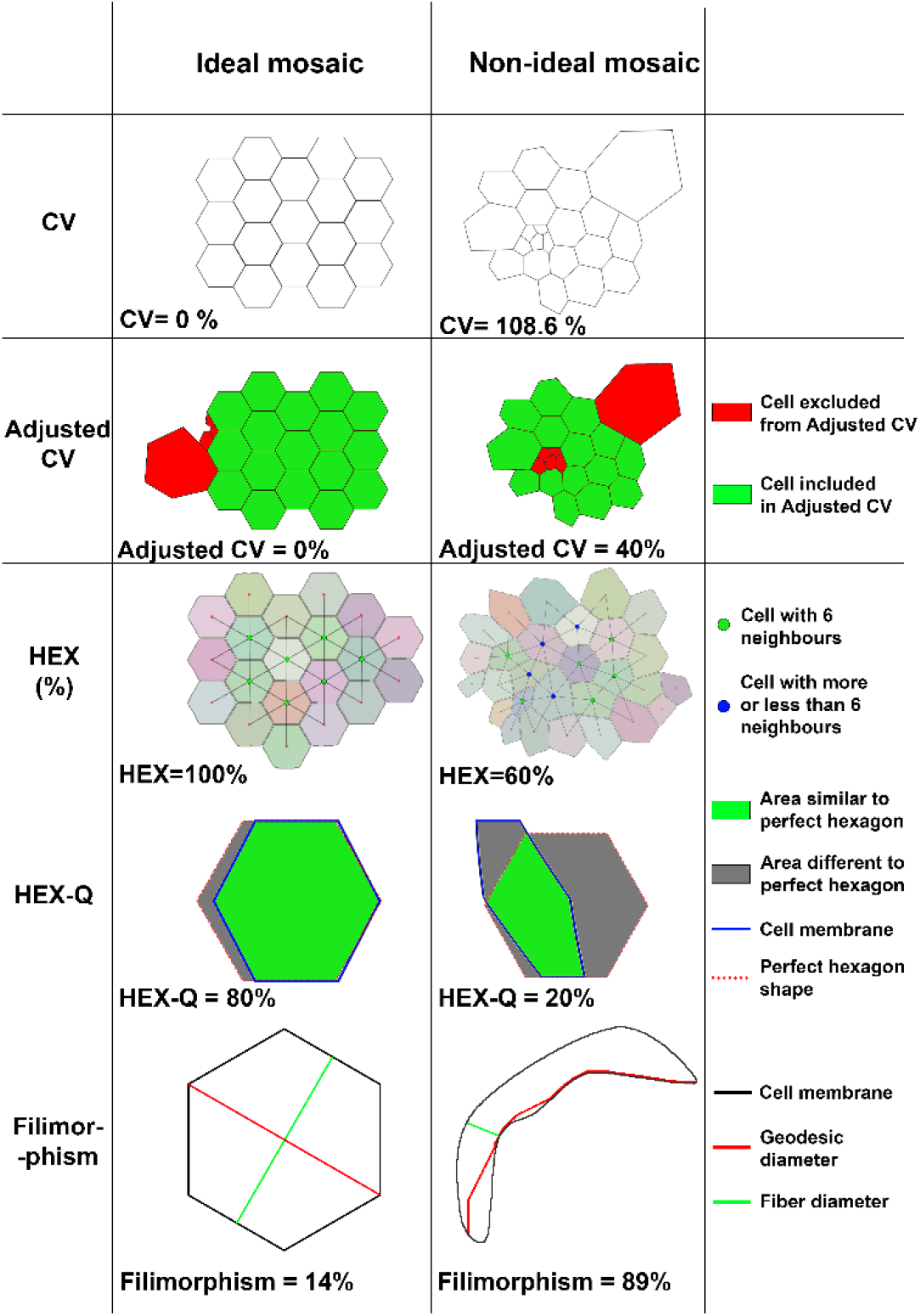
Parameters for quantifying hCECs morphology. CV: Coefficient of variation of cell area, HEX: percentage of hexagonal cells. Details for each parameter and their calculation method can be found in the supplementary methods.

1/ CV of cell area: the conventional parameter for polymegathism already mentioned. The lower the value, the more homogeneous the mosaic. It was calculated by the formula 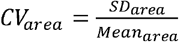. The CV of a healthy endothelium ranged from 26% to 35%, depending on the patient’s age [39], [40], [41], [42], [43]; 2/ Adjusted CV was specifically designed for this study to assess cell cultures containing several cell populations in terms of area. This adjusted CV allowed studying the CV of the majority population within the culture by replacing mean with the median and the standard deviation with another one calculated from median absolute deviation (see Supplementary methods); 3/ HEX for polymorphism. The HEX of an healthy endothelium ranged from 45 to 70% depending on the patient’s age [39], [40], [41], [42], [43]; 4/ Quality of hexagonality (HEX-Q) was also specifically designed for this study: among hexagonal cells (with 6 neighbors), it measured their proximity with a perfect hexagon based on HEXADEV criterion [44]. HEX-Q calculation took into account the convexity of the cell as well as the sides length and the angle of each side of the hexagonal cell (see Supplementary methods); 5/ Filimorphism: this parameter specifically designed for this study assessed cells elongation by modifying the aspect ratio formula (depending on definitions, aspect ratio in the literature is calculated either 1-as the ratio between the length of the major axis and the minor axis of the fitting ellipse of the cell or 2-as the ratio between the maximal and minimal Feret diameter of a cell) adapting it even for extremely concave (U-shaped) cells (see Supplementary methods).

### 2.9. Establishment of endothelial quality score

To comprehensively evaluate the different parameters, a score was established, hereafter referred as to the Endothelial Quality Score (EQS). This scoring system assigned 50% weight to ECD, as it is the primary criterion used in clinical practice (eye banking), and 50% to morphology, considering that both were equally important. Morphology evaluation was based on the five parameters previously described: HEX, HEX-Q, CV, adjusted CV and filimorphism, each contributing 1/10 of the EQS. The EQS was derived from Z-scores, which were calculated by the following formula: 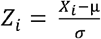 ; where Z_*i*_ was the score value, Xi the raw value, μ the population mean for the given criterion, and σ the standard deviation of the referred population. The higher the EQS, the better the endothelial quality.

### 2.10. Statistical analysis

To analyze results of the algorithm using the AI model, the software GraphPad Prism was used. Initially, the normality of each culture for each parameter was tested using three different tests (Kolmogorov-Smirnov test, Agostino and Pearson omnibus normality test, and Shapiro-Wilk normality test). If one of these three tests indicated normality for the four cultures, a Repeated Measures ANOVA followed by Bonferroni’s multiple comparison test was performed as a post-hoc analysis. If none of the normality tests showed normal distribution, a Friedman test was used, followed by Dunn’s multiple comparison post-hoc test.

To evaluate the relationship between coating molecules and ECD or morphology, a linear mixed effect model was used. ECD values and other positive criteria were log-transformed before the mixed effect analysis. This model accounted for both fixed effects, such as factors of interest (coating molecule), and random effects (donor, replicate ID), to account for inter-individual variations and correlations between observations. Statistical analyses were performed using R software (version 4.4.1), with the “lme4” and “lmerTest” packages to fit a linear mixed-effects model. Results were expressed as ratio of ECD effect relative to the reference molecule with the corresponding 95% confidence intervals.

## 3. Results

### 3.1. Efficiency validation of the morphological analysis

For the 4 cultures selected for validation, the 6 parameters successfully differentiated them as anticipated (Fig. 3). Culture 1 (EndoMT) had a low ECD with very low hexagonality and resembled fibroblasts, with high polymegathism. Culture 2 (double population) had an acceptable hexagonal pattern (HEX-Q) but significant pleomorphism (high CV and high filimorphism). For the last two cultures: culture 3 had a high ECD and low polymegathism; however, it exhibited higher pleomorphism than culture 4. Culture 4 had a slightly lower ECD, but his pleomorphism was minimal (HEX, HEX-Q, filimorphism).

**Fig. 3.**
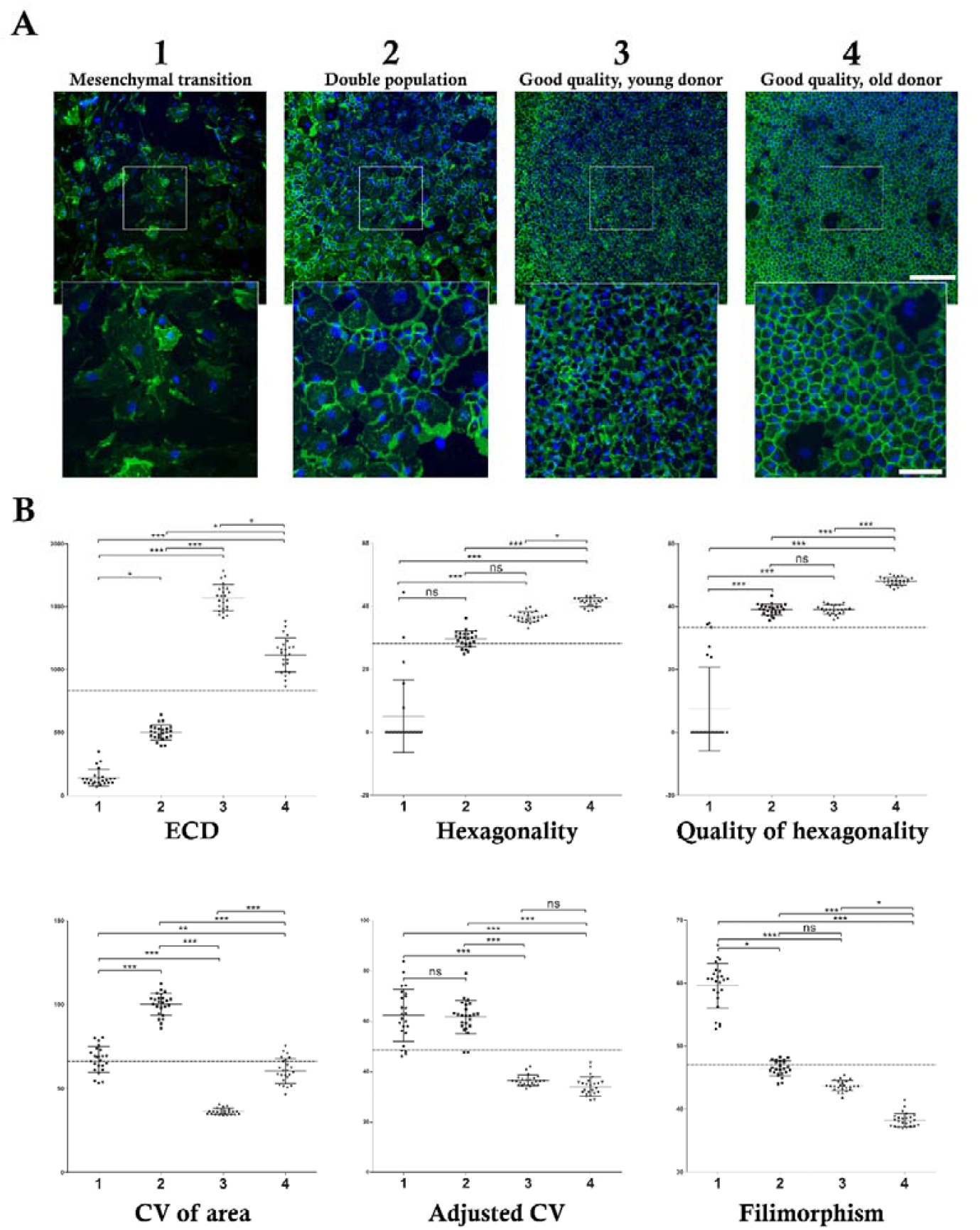
Validation of the AI-based method for quantifying the various hCECs morphological parameters. **(A)** Endothelial cell cultures of highly variable quality used for AI algorithm validation. Cell lateral membranes were labeled with anti-NCAM (in green) and the nuclei were counterstained using DAPI (in blue). The images were taken in the center of each well at x10 magnification with a zoom x6 in the center of the image. Image scale bar = 300 μm, zoom scale bar = 50 μm. **(B)** Evaluation of 6 parameters across 4 different culture qualities. Graph illustrating 6 objective evaluation parameters across 4 culture qualities: 1. EndoMT cells; 2. Mixed population quality; 3. High quality – young donor (below 30 yo), low passage; 4. High quality – aged donor (over 30 yo), high passage. ns=non-significant=p-value>0.05, *=p-value<0.05, **=p-value<0.01, ***=p-value<0.001).

The EQS significantly discriminated the two cultures with a good phenotype by assigning a significantly higher EQS to the culture with the highest ECD (culture 3), despite having slightly higher pleomorphism (Fig. 4). In contrast, the first two cultures were not significantly different. However, these two cultures appeared visually different with NCAM staining, with a complete absence of hexagonal-shaped cells resembling hCECs in culture 1. In culture 2, some cells exhibited a hexagonal shape similar to those observed in cultures 3 and 4.

**Fig. 4.**
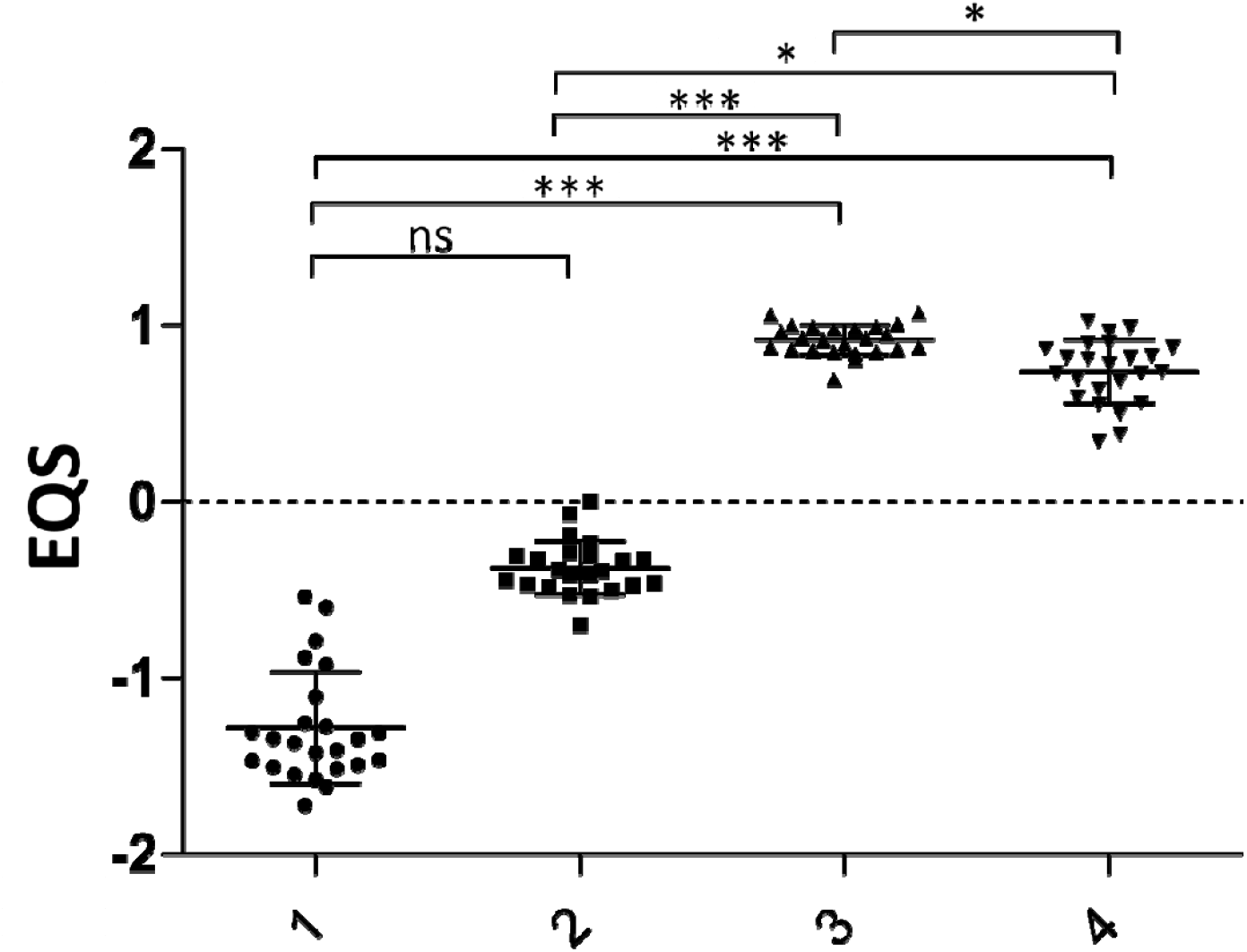
Endothelial quality score (EQS) for evaluating cell culture quality based on morphological and ECD criteria. EQS was calculated from the z-score of each of the 6 parameters chosen (ECD, Hexagonality, Quality of Hexagonality, CV, Adjusted CV, Filimorphism). *=p-value<0.05, **=p-value<0.01, ***=p-value<0.001).

### 3.2. Coating and donor impact on morphological and density parameters

Cell cultures from the 8 donors exhibited highly variable quality (Fig. S12 in the supplemental data), making it impossible to distinguish the effects of the different coatings by naked-eye observation and by conventional statistical analysis. The mixed model allowed us to assess their impact while considering multiple variables, including both intrinsic and extrinsic donor-related factors. An example of cell response to the different coatings was shown Fig. 5. For ECD, no molecule had a significantly lower value compared to the uncoated control, while height molecules exhibited significantly better results. For iMatrix-511, although ECD was not significantly higher than in the uncoated control, it demonstrated reduced cell size variability (adjusted CV), fewer filiform cells, and a higher proportion of cells appeared hexagonal in shape (HEX-Q). Only HEX did not allow for differentiating the different coating molecules. On the contrary, HEX-Q was able to discriminate between cultures (for instance B121, B332, and iMatrix-511 had higher HEX-Q than the control).

**Fig. 5.**
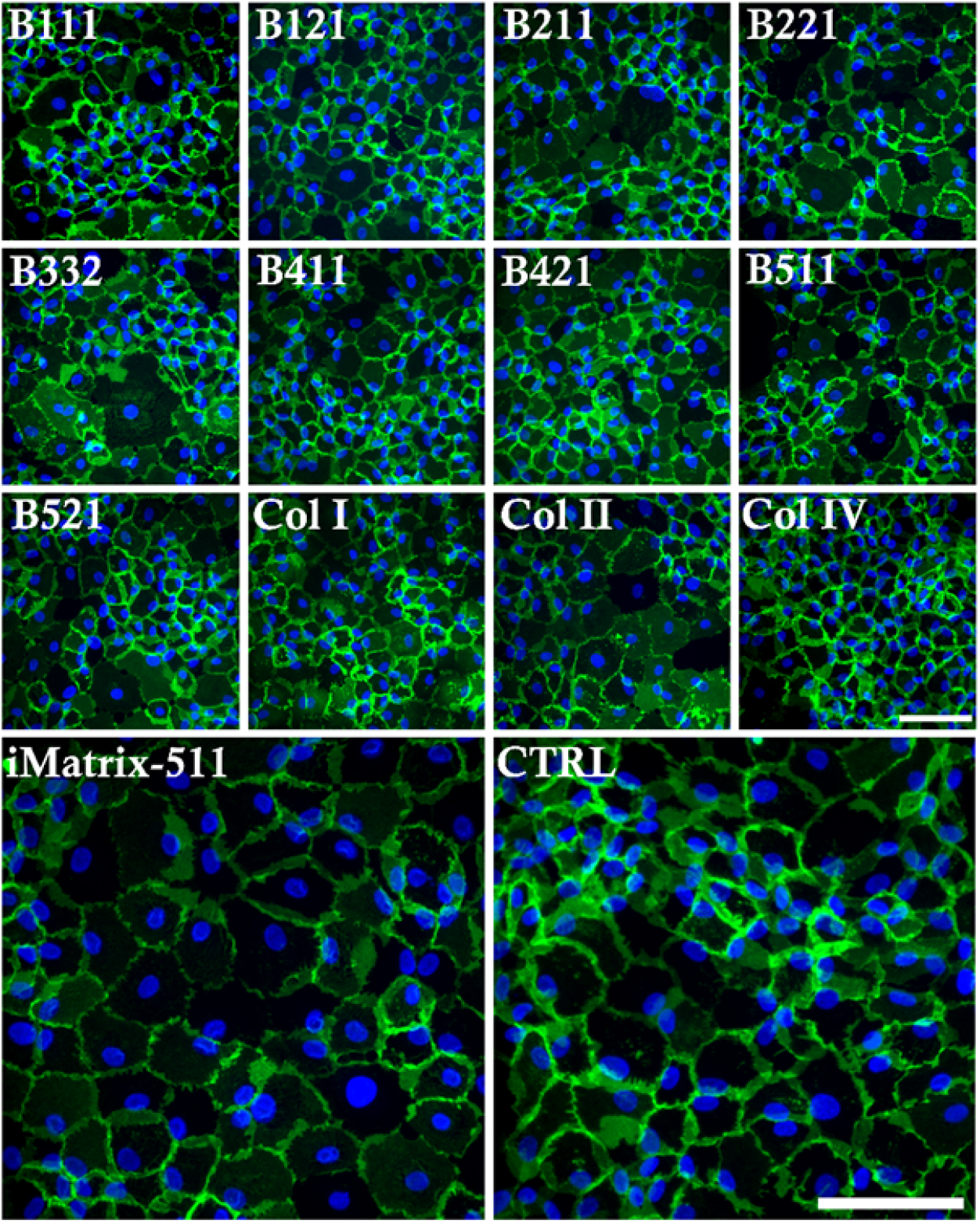
Representative example of the effects of coating molecules on hCECs. Cell lateral membranes were stained with anti-NCAM (green) and the nuclei were counterstained using DAPI (blue). Donor used for this example was “c” in Table 1. Scale bar 100 μm.

**Fig. 6.**
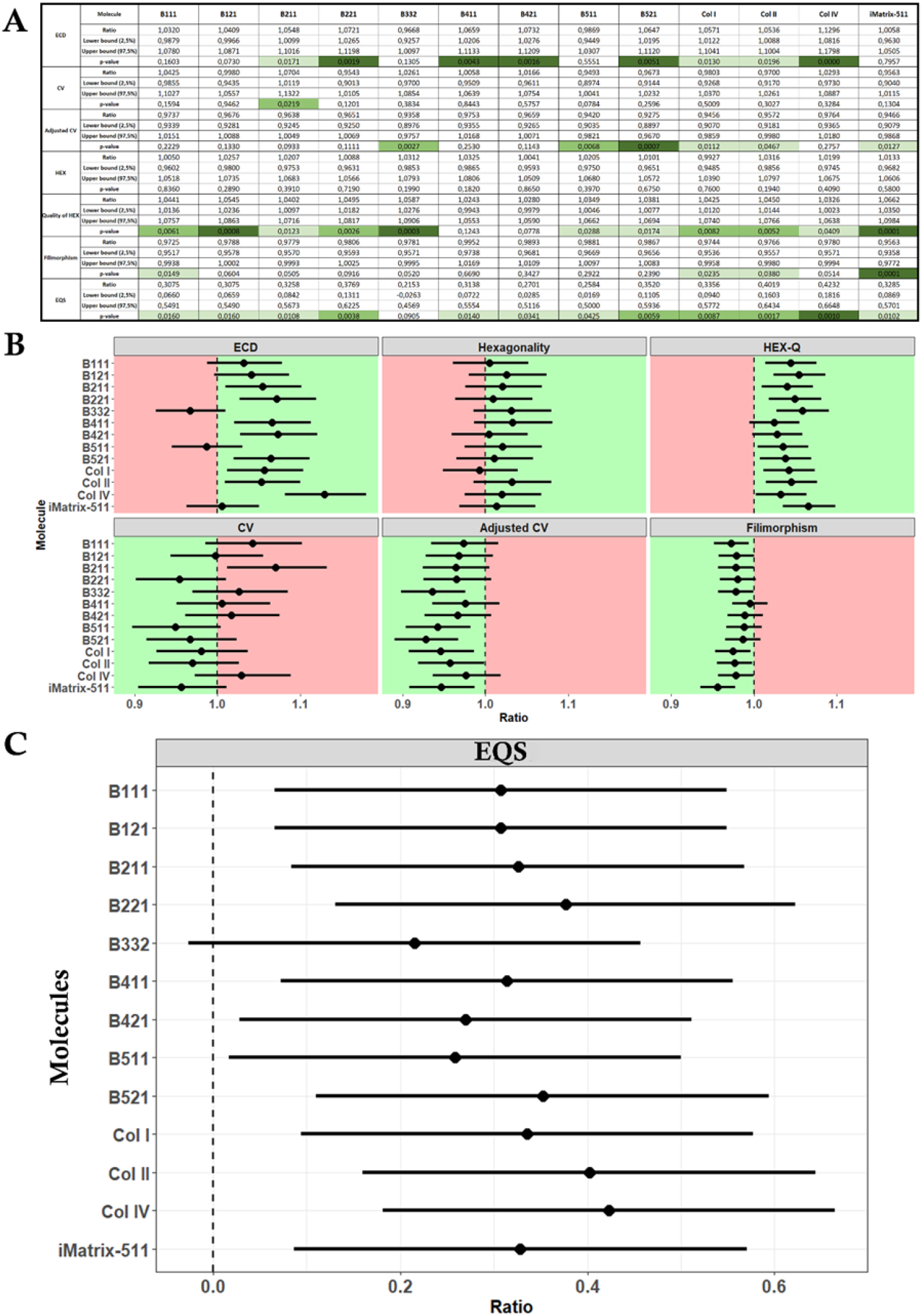
Summary of the mixed model analysis for coating molecules. **(A)** The table presented the ratio, p-value, and confidence interval, highlighting significant differences between the tested coating molecules and the control. The color scale indicates the level of significance, with darker green representing higher significance and lighter green representing lower significance. **(B)** Forest plot of molecule expression ratios according to the evaluated criterion. This forest plot illustrated the expression ratios of different coating molecules based on the evaluated criterion. Each point represents the mean ratio value, while the vertical bars indicate the corresponding confidence intervals (lower and upper bounds). The vertical line at y = 1 served as the reference value compared to the uncoated control. A ratio greater than 1 suggested superiority compared to the control, whereas a ratio lower than 1 indicated inferiority compared to the control. If the ratio of the parameters ECD, Hexagonality, and HEX-Q is greater than that of the control, the culture quality is better (green zone); otherwise, it is worse (red zone). If the ratio of the parameters CV, Adjusted CV, and Filimorphism is greater than that of the control, the culture quality is worse (red zone); otherwise, it is better (green zone). **(C)** Forest plot of molecule expression ratios (compared to the uncoated control) according to the endothelial quality score (EQS) (including all the 6 evaluations parameters). Each point represented the mean ratio value, while the horizontal bars indicated the corresponding confidence intervals (lower and upper bounds).

EQS differentiated the efficacy of coating molecules on cell culture quality. Only B332 did not have a significantly better EQS than the uncoated control, while all other coating molecules showed an improved one. The coating that achieved the highest score was collagen IV (Col IV). iMatrix-511 also improved cellular quality, but to a lesser extent.

## 4. Discussion

Corneal endothelial diseases are the leading indication for corneal transplantation worldwide. Two ATMPs are or will soon be able to compensate for the shortage of donors: cell therapy (available in Japan), and tissue therapy (in development) [45], [46]. To ensure their large-scale production, both techniques require the culture of hCECs of clinical grade, i.e. with the correct phenotype and obtained in large quantities according to pharmaceutical standards.

To draw a parallel with donor corneas, ECD is the main release criterion by corneal banks, and the decision threshold varies depending on the surgical technique considered. The threshold of 2000 cells/mm^2^ is consensual for penetrating grafts, and a higher value is required for endothelial grafts, as the surgery itself causes additional cell loss. Some banks add criteria for cell mortality, but these are not consensual. There are two historical morphometric parameters, used for 50 years to qualify the endothelium *in vivo* in patients and *ex vivo* in eye banks: the percentage of hexagonal cells (HEX) to evaluate polymorphism, and the coefficient of variation of cell surface (CV) for pleomorphism. HEX and CV are provided by eye banks but rarely serve as non-compliance criteria due to a lack of data on their postoperative impact [47]. Until recently, they were not used for the selection of *in vitro* cell cultures. Okumura et al. developed a software called the Corneal Endothelial Cell Analyzer to assess the transformation of hCECs into fibroblasts, based on cell density, percentage of hexagonal cells (HEX), and the variation coefficient based on the aspect ratio (ratio of height to width).

This software follows four processing steps: 1) noise reduction, 2) contour enhancement, 3) binarization, and 4) skeletonization [31].

Cellpose is an open-source software for segmenting cells in culture. Pretrained models available did not correspond to our needs, but the tuning of our own model was very efficient. The dissimilarity criterion c evaluated with a tolerance margin of 3 pixels (c = 0 in the ideal case and c > 1 for poor segmentation) was 0.034±0.010 when the model segmentation was compared to the ideal manual segmentation [38]. Additionally, we proposed several metrics to refine the assessment of hCECs cultures, as morphometric analysis (limited to HEX and CV) was initially developed for in vivo specular microscopy images, where endothelial cells exhibit a more regular morphology than in vitro: 1/the adjusted CV is more suitable than CV in the context of cultured hCEC likely to contain double or triple populations of normal, senescent and/or fibroblastit cells; 2/ HEX-Q is a sensitive criterion to evaluate the similarity of the cell contour to a regular hexagon; 3/ filimorphism, as a robust metric to study cell elongation. Further studies are now needed to establish relationship between our proposed metrics (considered as new image biomarker of cell quality) and conventional biological biomarkers already used in hCEC production.

To obtain a clinical-grade endothelial cell culture, several difficulties must be overcome. In some cultures, several different cell populations are observable, with varying phenotypes, ranging from typical endothelial cells to senescent or even fibroblastic cells (undergoing an EndoMT) [11], [14], [32], [48]. To approximate physiological conditions, the use of coating molecules on plastic culture plates is widely referenced [16], [20], [23], [25], [28], [30], [32]. The DM is composed of collagen types IV and VIII, laminin, fibronectin, and other extracellular matrix (ECM) proteins that provide structural support and influence cellular behavior [28], [49], [50], [51], [52], [53]. These proteins play a critical role in regulating fundamental cellular processes such as cell adherence, proliferation, polarity, morphogenesis, and function, particularly the integrin/laminin complexes [54]. Okumura et al. identified 2 laminin isoforms present in DM and demonstrated their positive effect, as well as that of the recombinant E8 fragment of laminin 511, on endothelial culture yield, in comparison with laminin 211, fibronectin, collagen I, FNC Coating mix (Athena Environmental Sciences, Inc., Baltimore, MD, USA) and no coating. Morphometric parameters were not used for their selection, which was based on adhesion tests, proliferation and flow cytometric analysis of endothelial marker expression [20], [55].

In our screening study, we compared a wide range of coating molecules: 9 laminins and 3 collagens selected from the known composition of DM [56], [57]. The use of NCAM labeling, employed for cell segmentation, relies on its ability to reveal lateral membrane expansions. Although ZO-1 is more commonly used, NCAM appear to be linked to activity and differentiation levels [58], [59]. Indeed, cultures 1 and 2 struggle to be statistically differentiated by our analysis, particularly because the cultures do not present hexagonal cells, giving two criteria with values close to zero and thus a score whose interpretation must be nuanced. The use of a segmentation based on two markers, one endothelial and one fibroblastic, could be interesting to complement the analysis and allow the discrimination of two cell population in the same culture. In that case, the AI will need to be retrained on this other marker.

To allow a global interpretation, we grouped the 6 morphometric parameters and DCE into an endothelial quality score, giving equal weight to both the quantity and quality of the cells. This score, which allowed the discrimination of cultures that were not very different, could be used during the production of endothelial ATMPs and compared to other critical quality attributes. The weighting of each parameter depends on the result of future studies (preclinical and clinical) can be adjusted if necessary.

One limitation of this work is the use of 384-well plates. While they offer the advantage of minimizing cell consumption, they are associated with greater inter-well variability due to their small volume, which amplifies the impact of slight variations in the number of seeded cells. For the qualification of cultures during production, larger surface areas will likely be more appropriate.

The robustness of our results relies on the use of several cultures with significant variability (extreme ages, passage, quality, initial DCE) and statistical analysis through a mixed model.

## 5. Conclusions

In conclusion, we demonstrated that our morphological analysis tool allows the discrimination of quality variations between very similar cultures. We then used this tool to identify a coating molecule capable of improving the *in vitro* culture of hCECs. To account for all the parameters of cell quality, we developed a score that provides an overall assessment of the culture quality, highlighting that the coating has a positive effect on the cultures, regardless of the laminin used. Using a complex statistical model, we were able to identify coating molecules that show better morphology or density compared to the uncoated control. Collagen IV appears to be the best coating molecule for improving hCECs culture in this context and, based on this analysis, even better than iMatrix-511.

## Acknowledgments

We extend our sincere gratitude to the cornea donors and their families for their generous and invaluable contributions to medical and scientific advancement. We also wish to sincerely thank the Eye Banks of Saint-Étienne, Besançon, and Nantes for their crucial support in providing corneas for research purposes. We also thank the transplant coordination of the CHU of Saint-Étienne and Paris Robert Debré as well as the laboratory of anatomy of the Faculty of Medicine of Saint-Étienne.

## Author contribution

G.Tr., L.C., Z.H., P.G., G.Th., C.M., took part in the concept and design of the study. G.Tr., L.C., I.A., C.M., G.B., E.O., C.M., performed experiments, collected and analyzed data. G.B., Y.G. and C.M., trained the AI. G.Tr., L.C., Z.H., E.O., P.G., G.Th., C.M. wrote-original draft preparation, G.Tr., L.C., Z.H., E.O., Y.G., A.M., P.G., G.Th., C.M. wrote-review and edited. All authors have read and agreed to the published version of the manuscript.

