## Supplementary material for "Investigating The Role Of Molecular Coating in Human Corneal Endothelial Cell Primary Culture using Artificial Intelligence-driven image analysis": Figure S12

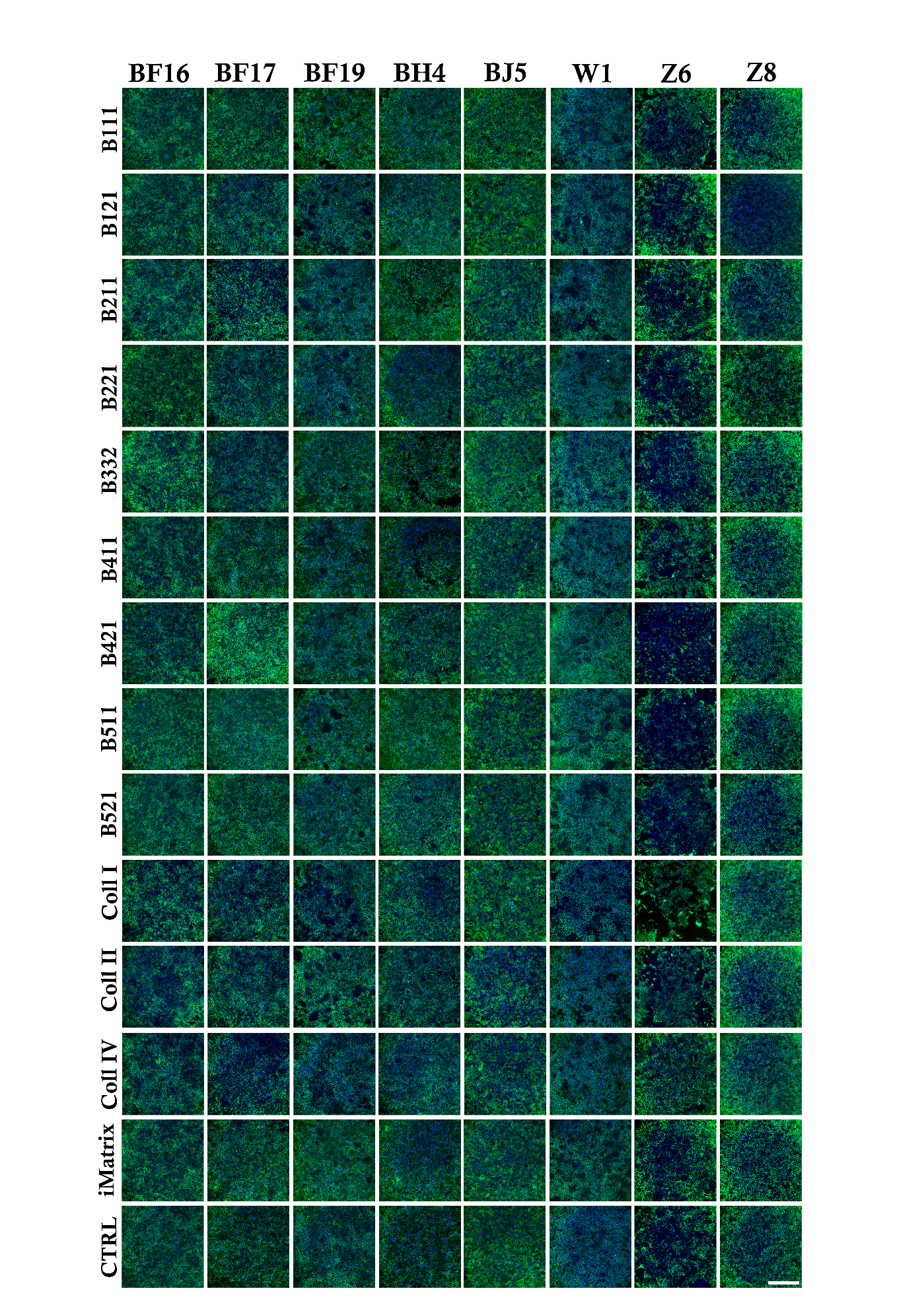

**Fig. S12.** Illustrative images of the 13 molecular coating compared to the uncoated condition (CTRL). Cell lateral membranes were stained in green by NCAM and the nuclei were counter-stained in blue using DAPI. The images were taken in the center of each well. The scale bar (on the bottom right image) was 500 µm for X10 images.
