## Supplementary material for "Investigating The Role Of Molecular Coating in Human Corneal Endothelial Cell Primary Culture using Artificial Intelligence-driven image analysis": Supplentary method

To evaluate the quality of the cultures, classical metrics such as ECD, polymegathism or coefficient of variation (CV), and pleomorphism or hexagonality (HEX) have already been developed [2]. The following text is describing method used to obtain those metrics in the present paper.

**Hexagonality**

Ideally, we aim for a maximum number of corneal endothelial cells with homogeneous area forming a regular hexagon with 6 neighbors. These metrics thus attempt to assess the resemblance to the ideal case:

$$\boldsymbol{H}\boldsymbol{EX}\left( \boldsymbol{\%} \right)\boldsymbol{=}\frac{\boldsymbol{n}_{\boldsymbol{HEX}}}{\boldsymbol{n}_{\boldsymbol{tot}}}\boldsymbol{\times}\boldsymbol{10}\boldsymbol{0}$$

Where $n_{HEX}$ is the number of cells with 6 neighbors in the image and *n_tot_* is the total number of cells.

Calculating hexagonality requires accessing neighborhood information per cell.

A direct approach might involve performing dilation cell by cell, taking the intersection with neighboring cells, and counting the number of non-zero components. However, this is a 2D approach that can be computationally expensive for a large number of cells. A faster approach might be to first calculate the contour of each cell, which is a very rapid operation using, for example, the opencv module [3].

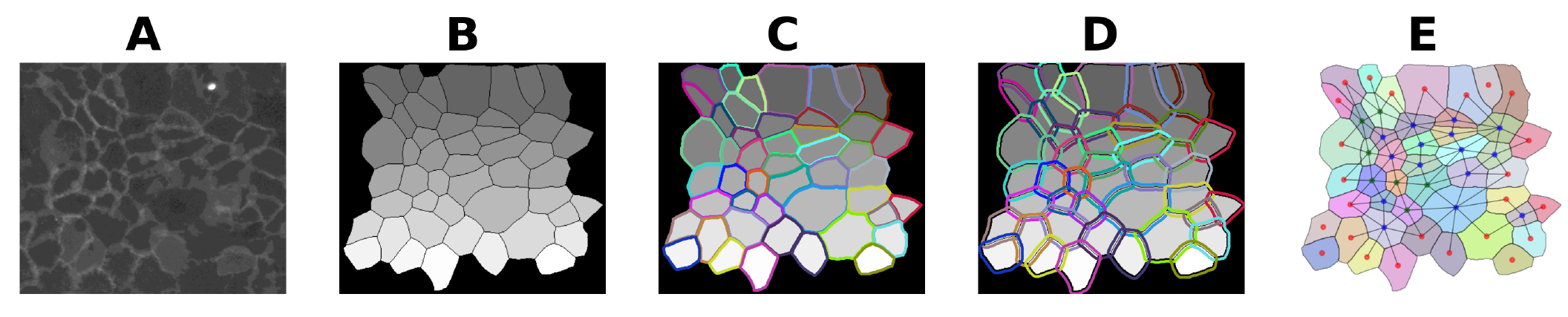

**Fig. S1.** Illustration of the method for calculating neighborhood information per cell. **(A)** An excerpt of an initial image of an hCECs culture labeled with NCAM. **(B)** Image labeling after segmentation by the Cellpose model retrained on our images. **(C)** Calculation of the contours of each cell. **(D)** Conversion of contours into polygons and dilation using the Python shapely module. **(E)** Calculation of neighborhood information with adjacency graph tracing. Red dot corresponds to the uncounted cells on the image edge, green dot are cells with 6 neighbours and blue dot are cells with more or less than 6 neighbours.

Next, we convert each of these contours into 1D polygons using the shapely module [4], as illustrated in C in Fig. S1. We then slightly dilate the polygons using the shapely module’s buffer option, as illustrated in D in Fig. S1. With these polygons, we perform intersections which are optimized operations in the shapely module, and we limit the number of intersections to calculate using bounding boxes of each polygon with an RTree structure [5]. These calculations have significantly accelerated the computations, with all operations (from contour calculation to polygon intersections) taking approximately 5 seconds for a 2048×2048 image with 2000 cells on our computer with NVIDIA RTX A4000 GPU and 11th Gen Intel(R) Core(TM) i7-11850H @ 2.50GHz CPU.

Finally, an illustration of the results is provided in E in Fig. S1. We then fill each contour with a randomly assigned color in slight transparency and trace a graph with nodes representing the centroids of each cell. The branches connect neighboring nodes. Some cells are on the periphery (i.e., not completely surrounded by other cells) and thus cannot be counted for the hexagonality calculation. Hence, they are marked with a red node. Cells with a green node have 6 neighbors (the ideal case) and are counted for the hexagonality calculation. Cells with a blue node are counted for the hexagonality calculation but do not have 6 neighbors. It should be noted that cells represented with a red node may be connected to cells marked with green or blue nodes, but two cells with a red node cannot be connected to each other as these do not count towards hexagonality. Moreover, this method can also be used to identify the triple points associated with each region. Indeed, triple points can be found in an image by convolving it with, for example, the filter: $\begin{matrix} 1 & 1 & 1 \\ 1 & 10 & 1 \\ 1 & 1 & 1 \end{matrix}$ on a binary skeleton image with non-zero pixels corresponding to cell boundaries connected in 8-connectivity. Then, we find values equal to 13 in the image. Once the coordinates of the triple points are calculated, we can use the previous polygons and the R-Tree structure to perform point-in polygon tests and thus find the triple points associated with each region in Python. This can be useful for calculating hexagonality with the vertex method quickly or for calculating HEXQ.

**Adjusted CV**

For the CV, we decided to work with a more robust definition, less sensitive to regions with very large areas corresponding to unanalysable acellular or senescent zones. We replaced the mean with the median and the standard deviation with a standard deviation calculated from the median absolute deviation (MAD) [6].

The current definition of CV is: $CV=\frac{\sigma}{\mu}$.

Using this metric involves modeling the distribution as a standard normal distribution $Z=\frac{X-\mu}{\sigma}$. We evaluated the relevance of using the traditional CV on the 15 labeled images from the database used for fine-tuning the Cellpose model. We illustrated a typical histogram of cell areas from one of these images in Fig. S2.

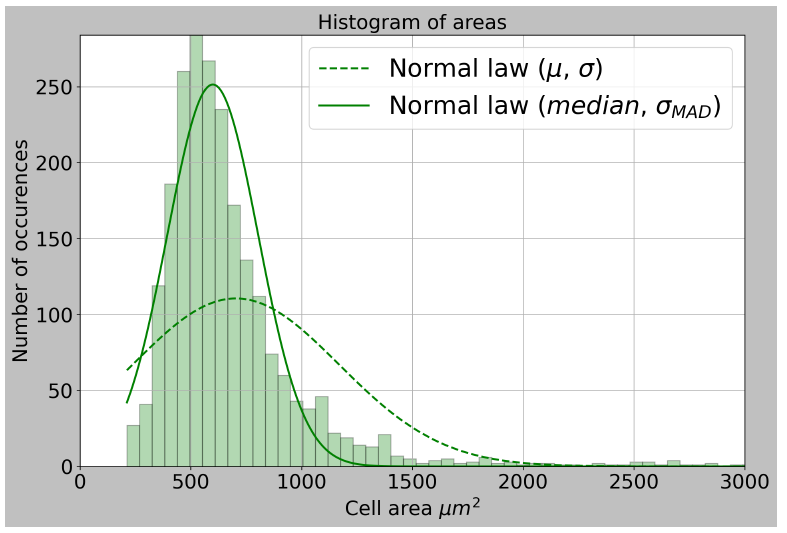

**Fig. S2.** Typical appearance of a histogram of cell areas from hCECs culture images labeled with NCAM. Example from one of our database images.

The model corresponding to the traditional CV is represented by a dashed line in Fig. S2. We can see that the traditional CV models the distribution of cell areas rather poorly. Next, we plot Henry’s lines, where the plotting method is described in [7], using the Benard method [8]. These lines provide richer information than the histogram. Indeed, some populations in the histogram are too minor and not visible. On Henry’s lines, the populations are more visible, and a population following a straight line can be modeled by a normal distribution.

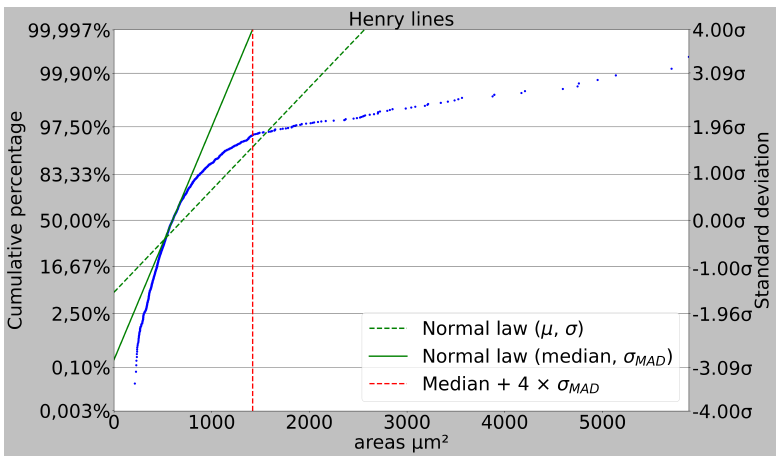

**Fig. S3.** Typical appearance of Henry’s lines of cell areas from hCECs culture images labeled with NCAM. Example from one of our database images.

Thus, in our case, as shown in Fig. S3, two main populations seem to be present: The first, corresponding to approximately 80 − 2.5 = 77.5% of the total population, is the majority population, that of cells with "normal" surface. The second population represents about 100 − 97.5 = 2.5% of the total population and probably corresponds to regions of very large areas associated with senescent or acellular zones. The remaining percentage of the total population corresponds to a "mixture" of the studied populations, i.e., areas that are not straight lines on Fig. S3. By plotting the straight line with the equation $Y=\frac{x-\mu}{\sigma}$ , represented by dashed lines in Fig. S3, we see that the current definition of CV does not model any of the studied populations. This can be explained by the fact that the mean and standard deviation used are too sensitive to outlier values, that is, values in very small proportions that deviate significantly from the majority values of the areas.

Our proposal is therefore to use a criterion based on the median, using the Median Absolute Deviation (MAD):

$$MAD=median(\left| x-median\left( x \right) \right|)$$

Where *x* is a vector *[area1, ..., arean]* containing all the areas of the studied cells. In the case of a normal distribution, we can write [6]:

$$\sigma_{MAD}=k\times MAD$$

*Equation 1*

where

$$k=\frac{1}{Ф^{-1}\left( \frac{3}{4} \right)}\approx\frac{1}{0.67449}$$

With Ф^-1^ being the inverse of the cumulative distribution function for the standard normal distribution and σ_MAD_ is the standard deviation calculated from the MAD. The use of MAD is not new, as it is often used to limit the influence of outliers in small proportions relative to the majority of the data; it is a robust measure useful in many contexts [6]. Thus, the new criterion we propose in our context is to evaluate "Adjusted polymegathism" (focusing on the main core of the distribution) with the following definition:

$$Adjusted CV= \frac{\sigma_{MAD}}{median(x)}$$

Which can be approximated by:

$$Adjusted CV\approx\frac{median\left| x-median(x) \right|}{median\left( x \right)\times0.67449}$$

where *σ_MAD_* is the standard deviation calculated according to equation 1 and *median(x)* is the median of the distribution of cell areas, with a ratio between the two quantities to have a dimensionless metric. Next, we plot on Fig. S3 the line with the equation $Y=\frac{x-median}{\sigma_{MAD}}$, represented by a solid line. We can see that we have managed to model the majority population quite satisfactorily, without considering acellular or senescent zones often corresponding to very large areas. Similarly, on the histogram shown in Fig. S3, the solid line represents the population model proposed by our updated CV, which also appears to model the studied population much better.

Thus, the previous definition of CV does not seem appropriate in our case. Indeed, the distribution can be modeled by a bimodal distribution (due to two populations), and the old CV tries to model it as a unimodal distribution. The old CV thus seems to work only when we have approximately one population under study. The new "coronal" CV proposed is more robust and allows us to satisfactorily model the majority population, without considering extreme values in very small proportions.

Finally, we focused on populations with areas between 0 μm² and *(median + 4×σ_MAD_)* μm². We chose *4σ_MAD_* to cover approximately 99.997 % of the majority population, beyond which the proportion of the studied majority population does not change significantly. Red dashed lines represent the value of *(median + 4×σ_MAD_)* on Fig. S3.

The cell areas did not pass the Shapiro-Wilk normality test (p-value at 0.05) between areas of 0 μm² and *(median + 4×σ_MAD_)* μm². Thus, we performed a non-parametric Mann-Whitney-Wilcoxon test to compare the distribution of the studied cells with that generated by a normal distribution with parameters *(median, σ_MAD_).*

The test was positive for 10 of the 15 images studied (p-value at 0.05), compared to 3 out of 15 images with a similar test for a normal distribution with parameters *(μ, σ)* over the interval 0 µm² and *(μ + 4×σ)* µm². Indeed, the population modeled by the Coronal CV may vary depending on the images; some images have cells with more heterogeneous shapes, limiting the extent of the Coronal CV modeling.

The only cases where the use of mean and standard deviation worked correspond to cases where there is approximately a single population of cell areas. However, for the majority of our images, the Coronal CV modeling is satisfactory and always models a significant proportion of the area population. This is not the case with the traditional CV, which almost never adequately models cell culture populations that almost always have more than one population of studied cell areas.

Moreover, we could also consider modeling the population with a more complex model composed of a mixture of normal distributions with different proportions. However, here we have limited ourselves to introducing the Adjusted CV, which is a simpler criterion more suited to our context than the traditional CV definition.

**Filimorphism**

For the analysis of human corneal endothelial cells, criteria for assessing "elongation" have already been studied [9]. This elongation can, for example, reflect the presence of asymmetry associated with the compression of hexagonal cells along their longitudinal axis. Therefore, several definitions of "elongation" already exist. For instance, we can mention the aspect ratio obtained by approximating the cell with an ellipse of the same second-order moment with the following formula [10]:

$$Aspect ratio=\left( 1-\frac{{Minor}_{axis}}{{Major}_{axis}} \right)\times100$$

or the aspect ratio formula as the ratio of the minimal Feret diameter to the maximal Feret diameter [9], [11]. Although these metrics can be effective in many cases, they may have certain limitations. Indeed, the most concrete example is the "saddle" morphology where the previous two definitions can induce errors. This example is shown, for instance, in (D) in Fig. S1. In practice, the two previous definitions of the aspect ratio are mostly useful in the case of convex cells or cells close to being convex. In this case, the approximation of elongation is quite good. This is the most common case for healthy or in vivo cells.

However, in the cases of cultures observed with NCAM imaging, cells can have very different morphologies from convex ones, and in fibroblastic cases with low-quality cells, they can "stretch" quite significantly in one direction compared to another. In this case, it may be useful to propose a more robust criterion characterizing the importance of this sometimes rather erratic elongation, where cells tend to resemble "threads".

Thus, we propose in this article the "filimorphism," a criterion that approaches 100% if the cell corresponds to a thread of infinite length or infinitesimal thickness, and that approaches 0% for a perfect disk, with dimensions equal in all directions. To achieve this, we used the definition of geodesic diameter proposed by Lantuéjoul et al. [12], and employed the method of calculating the fiber diameter proposed by Pourdeyhimi et al. [13].

**Geodesic Diameter**

To determine the length of a particle, we chose the definition of the geodesic diameter. To do this, we need to introduce the concept of a geodesic arc. Consider a cell *X* and two points *x* and *y* belonging to *X*. Several paths consisting of points belonging to *X* connect *x* and *y*, with the shortest one being called the geodesic arc, and its length is denoted *dX(x, y)*. The geodesic diameter is then defined as:

$$d_{geo}\left( X \right)={}_{x,y\in X}^{sup}{d_{X}(x,y)}$$

Its determination is illustrated in Fig. S4.

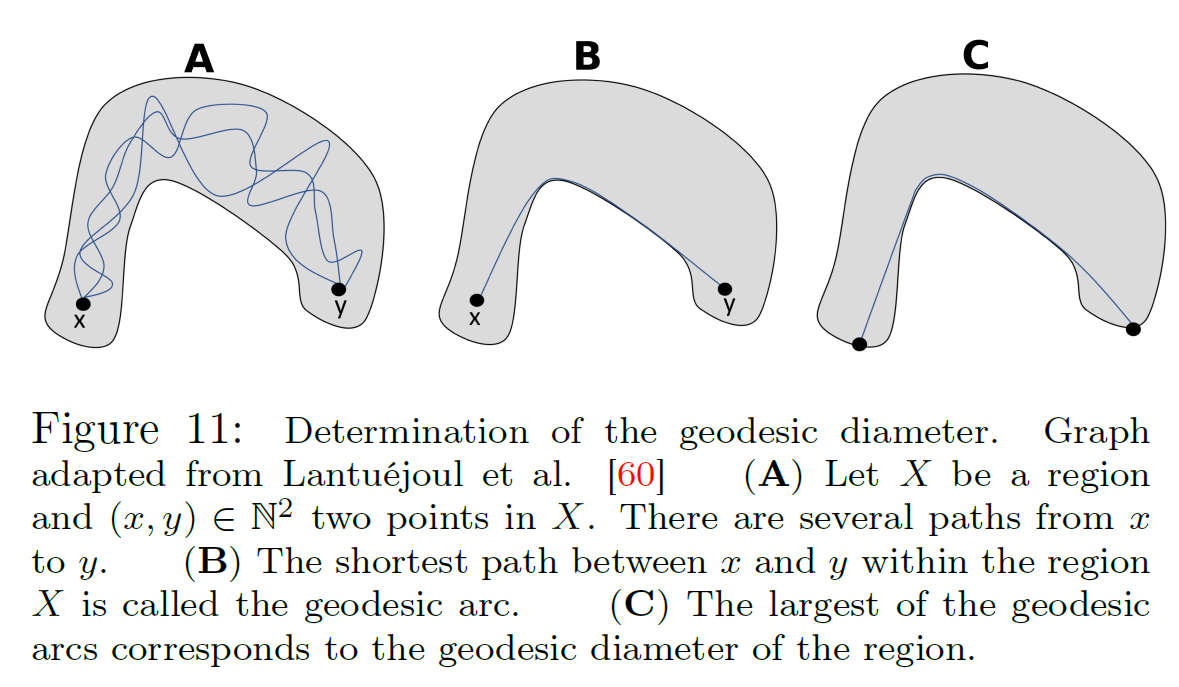

**Fig. S4.** Determination of the geodesic diameter. Graph adapted from Lantuéjoul et al. [12]. **(A)** Let X be a region and $(x,u)\in\mathbb{N}^{2}$two points in X. There are several paths from x to y. **(B)** The shortest path between x and y within the region X is called the geodesic arc. **(C)** The largest of the geodesic arcs corresponds to the geodesic diameter of the region.

To calculate it, we have proposed a Python code whose principle is detailed in Fig. S5.

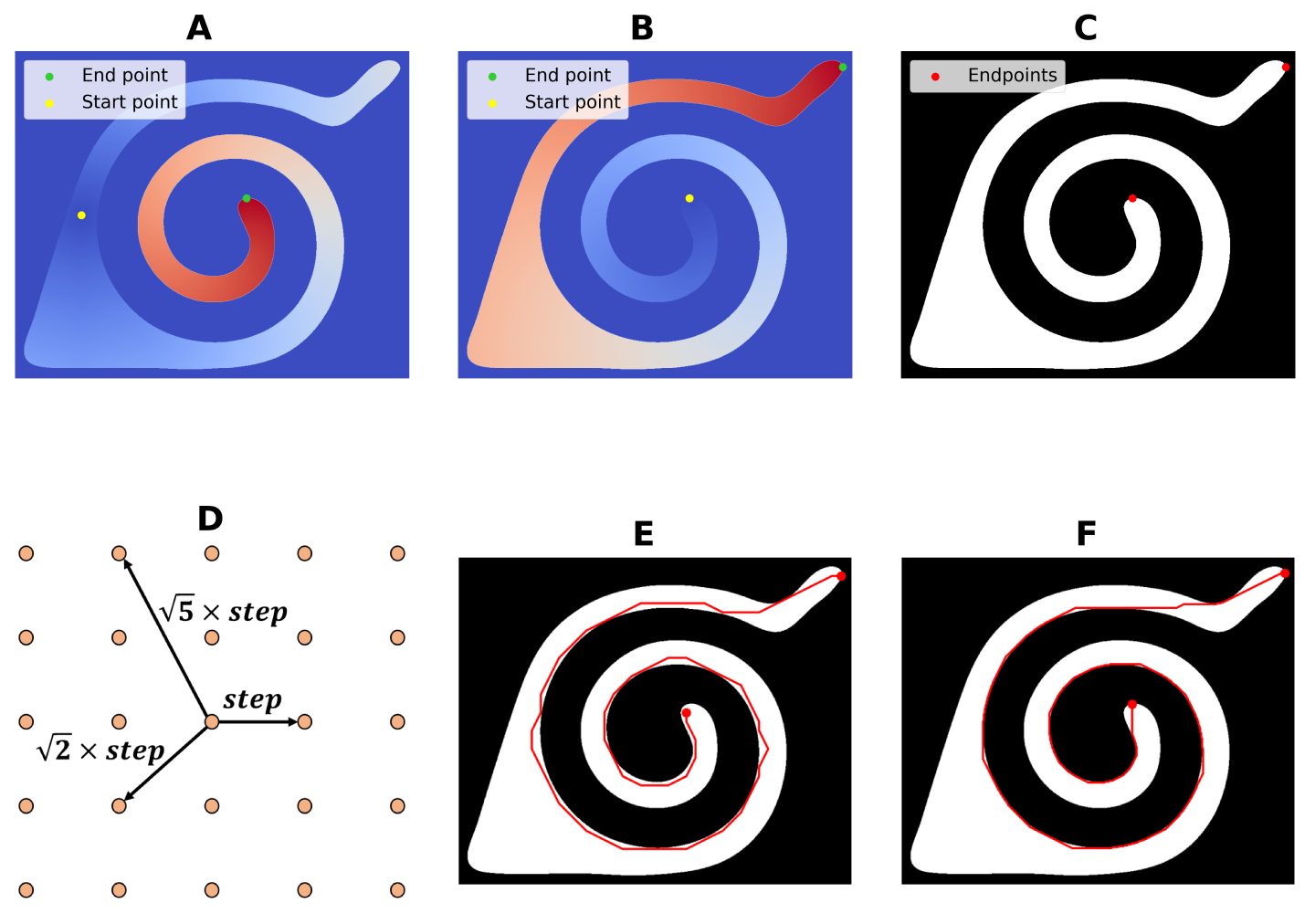

**Fig. S5.** Computational method for calculating the geodesic diameter. **(A)** Calculation of the propagation map following the various directions shown in **D** and starting from a point within the region (using the representative point function from shapely based on the region boundary polygon). **(B)** The maximum of the propagation map in **A** corresponds to the first point used to calculate the geodesic diameter. Calculation of a new propagation map from this point. **(C)** The maximum of the propagation map in **B** provides the second point used to calculate the geodesic diameter. The value of this pixel corresponds to the geodesic diameter. **(D)** It is also possible to plot the geodesic diameter within the region, for example by defining a graph with each node created from a 5 × 5 window that assigns weights corresponding to the distance associated with other nodes of the graph. Increasing the step value speeds up the calculations but reduces the accuracy of the results. **(E)** Application of Dijkstra’s algorithm to compute the shortest path between the two red points in **C** with the graph defined in the previous step with step = 20. **(F)** Application of Dijkstra’s algorithm to compute the shortest path between the two red points in C with the graph defined in the previous step with step = 2.

**Calculation of Fiber Diameter**

To determine the "fiber diameter," Pourdeyhimi et al.’s method [13] requires defining the concepts of morphological skeleton and distance map. A morphological skeleton is a "simplified" version of an image, with the possibility of reconstruction by an inverse operation to skeletonization [14], [15]. It involves taking the center of disks contained within the studied region with a maximal radius. Mathematically, the ball *Bn(x)* with center *x* and radius *n* is maximal with respect to the set *X* if there is no index *k* and center *y* such that:

$B_{k}(y)\subseteq B_{n}\left( x \right)$ *and* $B_{k}(y)\neq B_{n}(x)$

This means that there is no ball with center *y* and radius *k* that is strictly contained within *Bn(x).*

$$B_{n}(x)\subseteq B_{k}(y)\subseteq X$$

The morphological skeleton then corresponds to:

$$S\left( X \right)=\bigcup_{r} \in_{B_{r}(0)}(X)\setminus\gamma_{B_{1}\left( 0 \right)}(\in_{B_{r}\left( 0 \right)}\left( X \right))$$

where:

- $\in_{B_{r}\left( 0 \right)}(X)$ is the erosion of the set *X* by a ball of radius r.
- $\gamma_{B_{1}\left( 0 \right)}\left( \in_{B_{r}\left( 0 \right)}\left( X \right) \right)$ is the opening of the eroded set $\in_{B_{r}\left( 0 \right)}(X)$ by a ball of radius 1.
- $\bigcup_{r}$ denotes the union over all possible radii *r*.
- $\setminus$ denotes the set difference operation.

Next, the distance map of a binary image is defined using a given metric, in our case, the Euclidean metric [16]. Let *B* be a binary image. Let *deucl* be the Euclidean distance function and *B0* be the set of *(h, k)* such that *B[h, k] = 0*:

$$d(\left( i,j \right)B_{0}=min\left\{ \left. d_{eucl}(\left( i,j \right),(h,k) \right|(h,k)\in B_{0} \right\}$$

The method for calculating the fiber diameter is thus to perform the Hadamard product of the morphological skeleton image with the distance map [13]. Indeed, logically, since the distance map represents in a simplified manner the distance of each pixel in a region from the boundaries of the region, the "middle" value of the boundaries will approximately correspond to the Euclidean distance associated with the radius of the fiber under study.

Thus, it approximately corresponds to the center of the disks with a maximum radius fitting within the region. Since the skeleton can present branches according to the cell morphology that are too small to be representative of the actual morphology of the cell, the geodesic arc of the skeleton is kept by taking the closest starting point from the one used for the calculation of the geodesic diameter, and similarly for the endpoint.

This allows obtaining the approximate fiber diameter over the entire length of the studied cell without considering irregularities often related to the skeleton calculation, which is very sensitive to noise in the image. The search for the closest point can be optimized using, for example, a Kd-tree structure [17]. This selection of the geodesic arc between these two points is illustrated between B and C in Fig. S5.

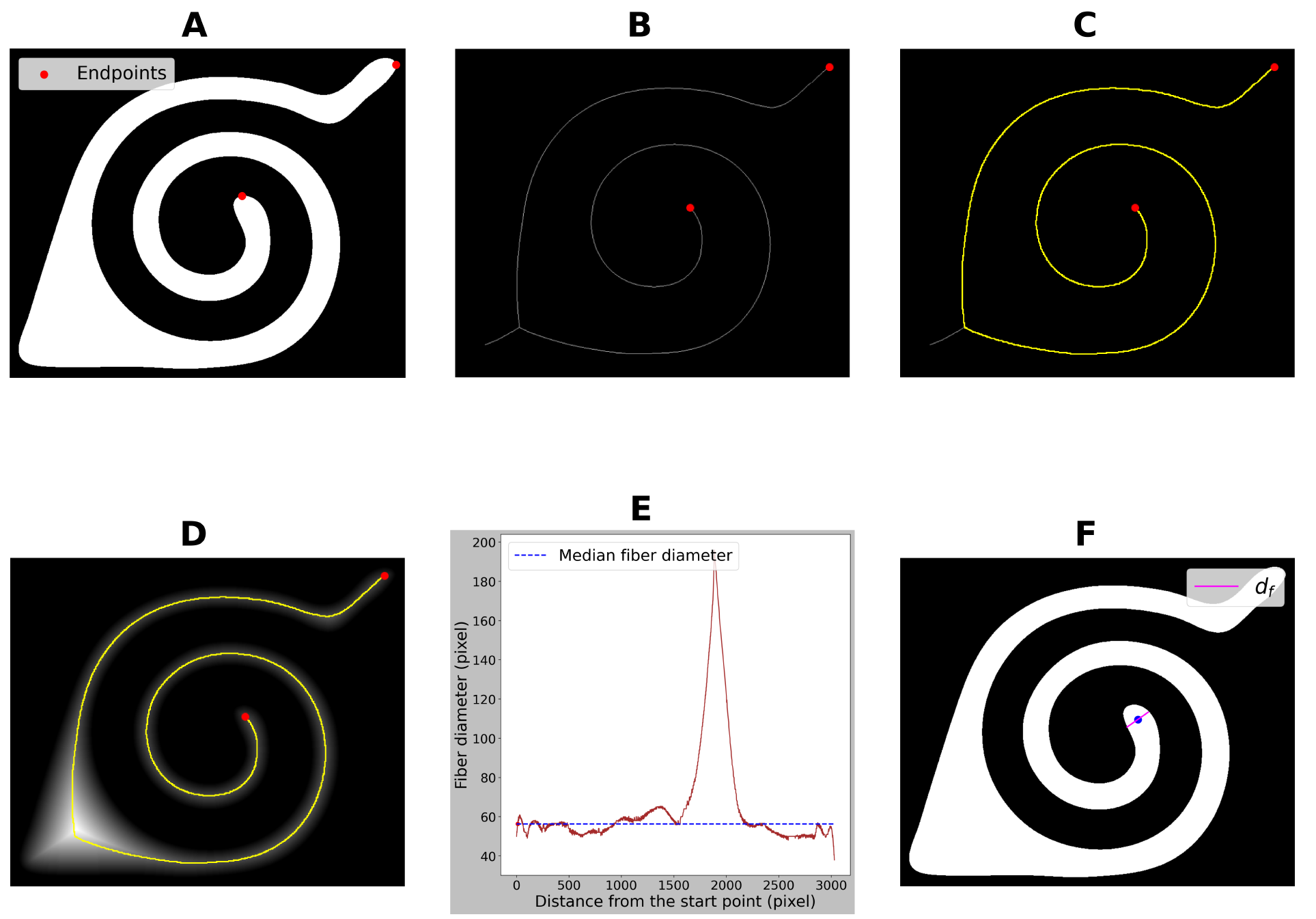

**Fig. S6.** Computational method for calculating the median fiber diameter. **(A)** Pair of points used to calculate the geodesic diameter of the region, determined in **C** in **Fig. S5**. **(B)** Determination of the points belonging to the morphological skeleton of the region that are closest to the points shown in the previous step using a Kd-tree structure [17]. **(C)** Calculation of the shortest path between the two points from the previous step using Dijkstra’s algorithm. **(D)** Multiplication of the previous part of the morphological skeleton by the distance map. **(E)** Evaluation of the value of the distance map along the yellow line from the previous step starting from one of the red points to the second. A peak is logically observed when the diameter of the figure is larger. To be robust against potential local irregularities in the region, we calculate the median of the calculated diameters. **(F)** Illustration in pink of the median fiber diameter value at one of the points on the distance map equal to the median fiber diameter. This corresponds to one of the intersection points of the red curve in the previous step with the blue dashed line.

Let $n,m\mathbb{\in N,}I\epsilon\mathbb{M}_{,nm}\left( \left\{ 0,1 \right\} \right)$, we assume that *I* contains only one non-zero region represented by a set of connected pixels with a certain connectivity of intensity 1. *Iskel* is the image corresponding to the morphological skeleton of *I* where only the geodesic arc between the closest skeleton points to those used for the calculation of the geodesic diameter is retained. *Iedt* is the distance map of *I* using an Euclidean metric. Inspired by the method of Pourdeyhimi et al. [13], we can then write:

$$I_{diameter}=I_{skel}\odot I_{edt}$$

Where ⊙ denotes the Hadamard product between two matrices.

We can then set *xstart* as the starting point used for the calculation of the previous geodesic arc. Since the skeleton is composed, after previous treatments, of a single 1-pixel wide line where each pixel is connected with 8-connectivity, there exists only one possible path in 8-connectivity between any non-zero pixel *x* in *Iskel* and *xstart*.

Let $X_{skel}\mathbb{\subset R^{2}}$ be the set of coordinates of the non-zero pixels of *Iskel*, $I_{skel} , \forall x\in X_{skel} , d(x,x_{start})$ is the length of the path between *x* and *xstart*. By plotting the curve, $\forall\left( i,j \right)\in X_{skel} , I_{diameter}\left[ i,j \right]=f(d\left[ i,j \right], x_{start})$, as illustrated in E in Fig. S6, we can notice that the two ends of the curve correspond to minimum values. This is verified in most cases, as it is the proximity to the region boundaries at the ends where the distance map logically has a lower value.

To avoid considering this edge proximity effect, the working interval is chosen between the first local maximum starting from the right of the curve and the first from the left. Then, if the two local maxima (from the left or the right) are equal, the maximum is global and the fiber diameter *df* is then taken as equal to the global maximum. Otherwise, it is taken as equal to the median of the curve between the two previous local maxima. The median is chosen instead of the mean to have a fiber diameter value less sensitive to potential irregularities along the particle. The process of calculating *df* is illustrated in Fig. S6.

**Definition of Filimorphism**

Therefore, the "filimorphism" proposed, characterizing the resemblance of a cell to a fiber, is defined as:

$$Filimorphism \left( \% \right)=\left( 1-\frac{d_{f}(X)}{d_{geo}(X)} \right)\times100$$

Filimorphism has the advantage, as illustrated in Fig. S7, of being more robust in cases of saddle-shaped.

or non-convex cells, which can be the case in cell culture or NCAM imaging, for example. It provides values close to aspect ratios with ellipse-related or Feret diameter-related definitions in the case of more conventional cell shapes. It should be noted that the aspect ratio definition with ellipses does not distinguish between hexagons and disks, unlike the Feret diameter or filimorphism definitions.

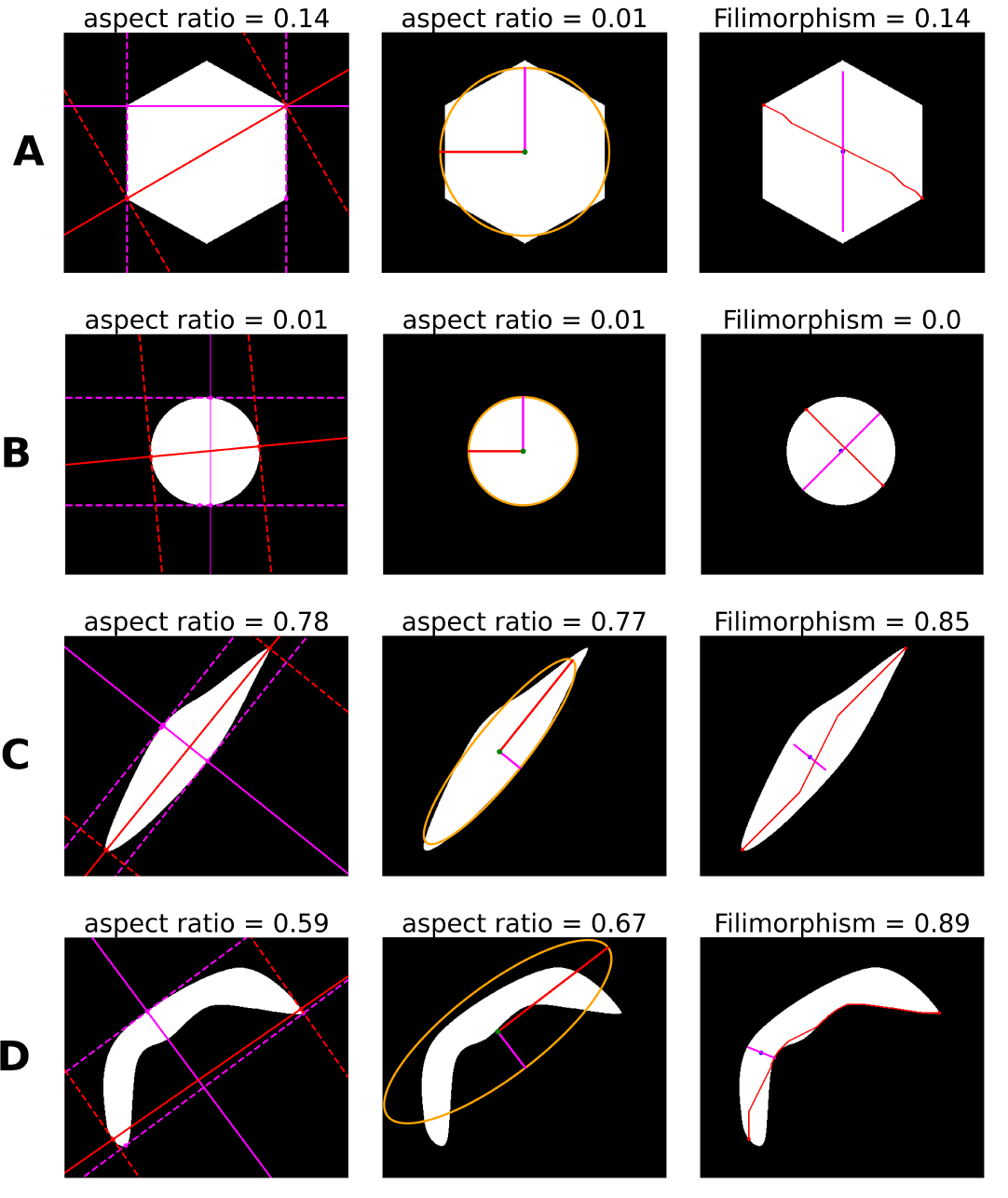

**Fig. S7.** Comparison between filimorphism and classical definitions of aspect ratio. The first column corresponds to the method using the Feret diameter, the second to the ellipse method, and the third column corresponds to filimorphism **(A)** Regular hexagon **(B)** Disk **(C)** Convex elongated cell (**D**) Example of an elongated "saddle" cell with a low degree of convexity.

**Hexagonality Quality (HEX-Q)**

In the ideal case, an endothelial cell is a regular hexagon with a relatively small area (on average 320 μm² for healthy men between 20 and 50 years old [18]). In the literature, hexagonality (i.e., the percentage of cells with 6 neighbors) is widely used to study the morphology of the corneal endothelium.

However, this criterion does not provide additional information on the similarity of a hexagon to a regular hexagon. Indeed, in the ideal case, endothelial cells are arranged in a regular hexagonal tiling to minimize the surface tension energy. As previously presented, a HEXADEV criterion has been published in the literature to study the similarity of a hexagon to a regular hexagon [19], defined as *(((A/P2)/0.07217) × 100) − 100(%)*. In our context, we have tested this parameter and proposed another criterion to make it more sensitive.

Therefore, we propose a criterion tending towards 100 for a regular hexagon and 0 if the morphology of the hexagon deviates too much from a regular hexagon.

Thus, we started from the definition of a regular hexagon. It is a convex polygon with 6 sides, all of equal length, and with all internal angles equal to 120°. We used a function with 3 criteria: one characterizing the convexity of the polygon formed by the cell, one the homogeneity of the side lengths, and the last the homogeneity of the internal angles.

**Convexity Criterion**

For the convexity criterion, the first step is to determine the positions of the triple points of the region (to study the polygon formed by connecting the pairs of consecutive points among the 6 triple points of a hexagon). To do this, starting from a segmentation corresponding to a binary image with boundaries represented by pixels of intensity 1 connected with 8-connectivity and the rest at 0.

With boundaries being single-pixel width lines, a quick calculation is possible. For example, a convolution with the filter $\begin{matrix} 1 & 1 & 1 \\ 1 & 10 & 1 \\ 1 & 1 & 1 \end{matrix}$ allows the determination of the positions of the triple points by identifying pixels with an intensity of 10 + 3 = 13. It is important to ensure that the central pixel of the filter is greater than or equal to 6, otherwise, other pixels than the triple points may have intensities equal to the sum of the central pixel intensity of the filter and the value three.

Once the coordinates of the triple points associated with the studied region are determined, we can study the polygon formed by connecting pairs of consecutive points among the 6 triple points of a hexagon.

Let $n,m\mathbb{\in N,}I\in\mathbb{M}_{n,m}\left( \left⟦ 0,1 \right⟧ \right)$ , assuming that *I* contains only one non-zero region represented by a set of connected pixels with a certain connectivity of intensity 1.

Let $X\mathbb{\subset R^{2}}$ be the set of coordinates of the non-zero pixels of *I*.

Let *(p1, . . . , p6)* be the coordinates of the triple points associated with the region, sorted based on the angle formed by the vectors $\vec{O_{x}}$ and $\vec{P_{c}P_{k}}$ for all $k\in\left⟦ 1,6 \right⟧$, where $\vec{O_{x}}$ is the horizontal unit vector of the coordinate system. *Xp* corresponds to the set of coordinates of the filled polygon formed by the points *(p1, . . . , p6)*. *Xc* corresponds to the set of coordinates of the pixels forming the convex hull of the points *(p1, . . . , p6)*.

Let *XΔXp = (X ∪ Xp) \ (X ∩ Xp)* be the symmetric difference between *X* and *Xp*:

$$C_{convex}=\left\{ \begin{aligned} \left( 1-\frac{\#(X\Delta X_{p})}{\#X_{p}} \right)\frac{\#X_{p}}{\#X_{c}} if {\#X}_{p}>{\#(X\Delta X}_{p}) \# \\ 0 otherwise \end{aligned} \right.$$

\# denotes the cardinal operator (i.e., the number of pixels within the studied domain). An illustration of *XΔXp* is provided in E on Fig. S8.

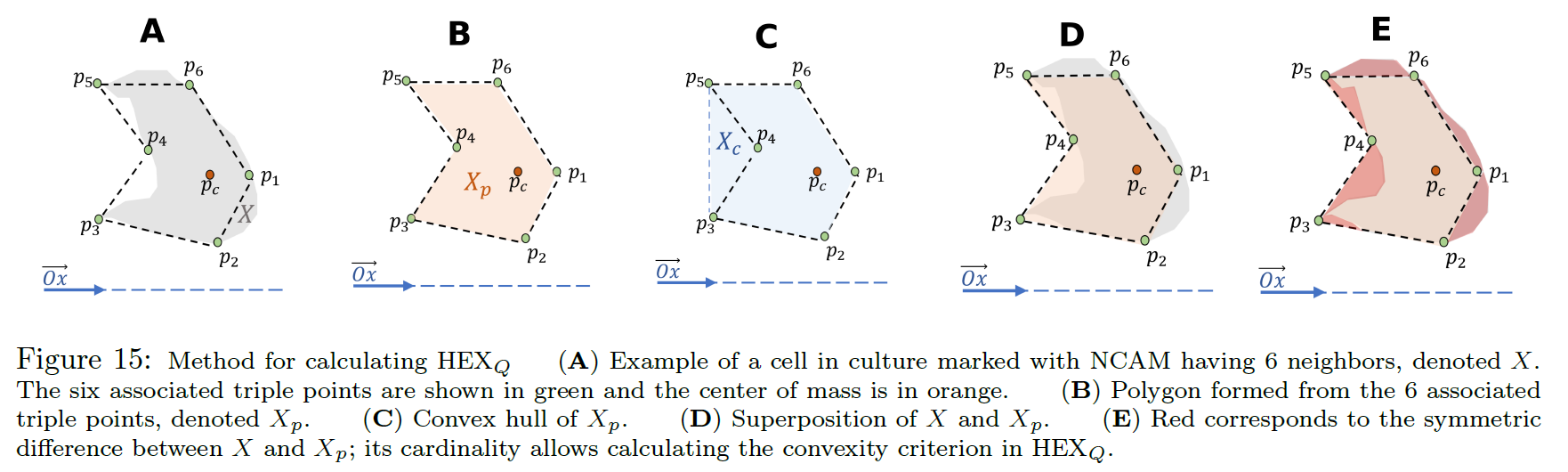

**Fig. S8.** Method for calculating HEX_Q_. **(A)** Example of a cell in culture marked with NCAM having 6 neighbors, denoted X. The six associated triple points are shown in green and the center of mass is in orange. **(B)** Polygon formed from the 6 associated triple points, denoted Xp. **(C)** Convex hull of Xp. **(D)** Superposition of X and Xp. **(E)** Red corresponds to the symmetric difference between X and Xp; its cardinality allows calculating the convexity criterion in HEX_Q_.

**Side Length Criterion**

Let *(p1, … , p6)* be the triple points associated with the studied region, sorted based on the angle formed by the vectors $\vec{O_{x}}$ and $\vec{P_{c}P_{k}}$ with $k\in\left⟦ 1,6 \right⟧ ,$ where $\vec{O_{x}}$ is the horizontal unit vector of the coordinate system.

Define:

$$\left\{ \begin{aligned} d_{i}=\sqrt{\left( x_{i}+1-x_{i} \right)^{2}+(y_{i}+1-y_{i})^{2}} \\ d_{6}=\sqrt{\left( x_{6}-x_{1} \right)^{2}+(y_{6}-y_{1})^{2}} \end{aligned}\forall i\in\left⟦ 1,5 \right⟧ \right.$$

where for all $\left( i,j \right)\in\left⟦ 1,6 \right⟧\times\left⟦ 1,6 \right⟧,(x_{i},y_{i}\mathbb{)\in N^{2}}$ are the coordinates (or spatial positions) of the point *pi*. *di* represents the Euclidean distance between two consecutive triple points *pi* and *pi+1*.

$${CV}_{length}=1-\frac{\sigma(\left[ d_{1},\ldots,d_{6} \right])}{\mu(\left[ d_{1},\ldots,d_{6} \right])}$$

where *σ* denotes the standard deviation of the vector and *μ* denotes the mean of the vector. We set *CVlength* =

0 if *σ* > *μ*.

**Internal Angles Criterion**

Using the previous notations, define:

$$\left\{ \begin{aligned} a_{i}=∡p_{k}p_{k+1}p_{k+2} \\ a_{5}={∡p}_{5}p_{6}p_{1} \forall_{i}\in\left⟦ 1,4 \right⟧ \\ a_{6}=∡p_{6}p_{1}p_{2} \end{aligned} \right.$$

where ∡*(ABC)* denotes the angle formed by the points *A*, *B*, and *C*.

$${CV}_{angle}=1-\frac{\sigma\left( \left[ a_{1},\ldots,a_{6} \right] \right)}{\mu\left( \left[ a_{1},\ldots,a_{6} \right] \right)}$$

where σ denotes the standard deviation of the vector and μ denotes the mean of the vector. We set *CVangle* = 0 if *σ* > *μ*.

**Definition of HEXQ**

For a region *X*, associated with 6 triple points *(p1, … , p6)*:

$${HEX}_{Q}\left( \% \right)=\left( C_{convex}^{\alpha}\times C_{length}^{\beta}\times C_{angle}^{\gamma} \right)\times100$$

with $\left( \alpha,\beta,\gamma\right)\in\mathbb{R}^{3}$ being coefficients that can be chosen empirically to better distinguish between good and poor cases. We propose *α* = 1, *β* = 1.3 and *γ* = 0.6.

**Tests and Comparisons with HEXADEV**

To compare the different metrics, we drew inspiration from the method proposed in the article [20]. We performed morphological variations of varying degrees and compared the values of the different metrics. For HEXADEV, the area and perimeter are calculated as follows in the original article [19]:

**Area (Shoelace Formula):**

$$\boldsymbol{A}=\left| \sum_{i=2}^{n+1} \frac{1}{2}\times(\left( x_{i-1}\times y_{i} \right)-\left( x_{i}\times y_{i-1} \right)) \right|$$

***Equation 2***

**Perimeter**:

$$p=\sum_{i=2}^{n+1} \sqrt{\left( x_{i}-x_{i-1} \right)^{2}+\left( y_{i}-y_{i-1} \right)^{2}}$$

where *xi* and *yi* are the coordinates of the *i*-th triple point and *n* is the number of triple points, with *(x1, y1) = (xn+1, yn+1)*. The problem with using these formulas, as discussed in [19], is that they consider the distance between consecutive triple points as a straight line, without accounting for potential tortuosity. Using these formulas, we will refer to the calculation of HEXADEV via the coordinates of the 6 triple points as "HEXADEV triple points".

To generalize this definition, we can propose calculating HEXADEV by determining the area through the sum of pixels of the studied region. We can also compute the perimeter using the Crofton perimeter (for example, this perimeter is presented in [21]). We will refer to this second method as "HEXADEV pixels".

Finally, in the following figures, we compare the different metrics across several types of cell morphology. The position of the triple points was automatically determined by a convolution as previously described, after positioning 6 hexagons identical to the studied hexagon around it.

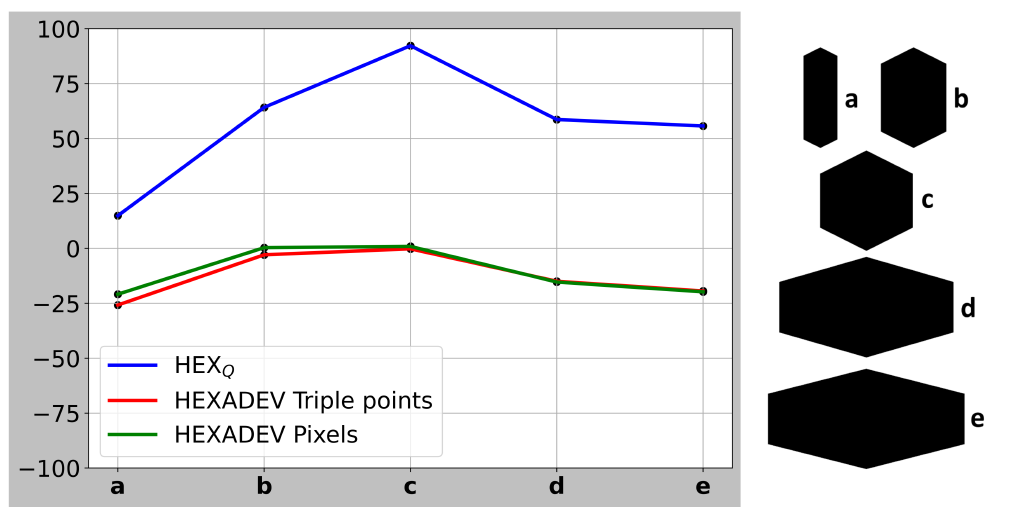

**Fig. S9.** Comparison of metrics assessing the similarity of a hexagon to a regular hexagon. Variation of the distance between two parallel sides of a hexagon.

By examining Fig. S9, where we progressively increase the distance between two parallel sides of a hexagon, we see that the three criteria seem to follow similar trends. However, HEXQ appears to exhibit more significant variations between different morphologies, notably between shapes b and c, which might allow for better differentiation. We can observe that HEXQ is not exactly equal to 100% for shape c, which seems to resemble a perfect regular hexagon. In reality, the position of the triple points may be very slightly altered from the ideal positions, and the criterion related to convexity might be slightly modified from 100% due to image resolution and minor pixelation effects. A value above 90% for HEXQ already indicates a hexagon very close to a regular hexagon.

Next, in Fig. S10, where we vary the degree of convexity of the hexagon, we see a limitation of the HEXADEV Triple points criterion. This criterion calculates the perimeter and area solely from the coordinates of the 6 triple points. Thus, if the triple points change little or not at all, as is the case here since the 6 neighboring hexagons do not change position, the value calculated by HEXADEV Triple points does not vary much. The degree of convexity of the hexagon is therefore almost not accounted for by this criterion. This is where HEXADEV pixels comes into play, allowing us to follow approximately the same trends as HEXQ between shapes b to e. HEXQ seems to better distinguish shapes a and b.

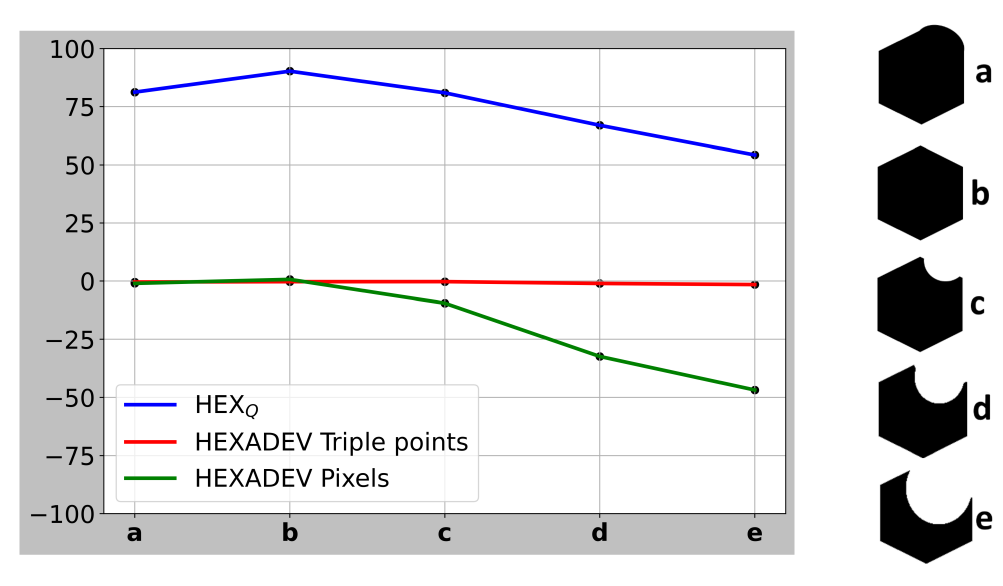

**Fig. S10.** Comparison of metrics assessing the similarity of a hexagon to a regular hexagon. Progressive change in the degree of convexity.

Finally, Fig. S11 shows the effect of simultaneously increasing the length of two parallel sides of a hexagon. Here, the HEXADEV pixels criterion shows very slight variations, and distinguishing the different shapes appears rather difficult. This situation is likely caused by approximations due to pixelation in the calculation of the perimeter and area. For the HEXADEV Triple points calculation, as the position of the triple points changes, the potential limitations presented in Fig. S10 are no longer present, and the curve shows variations similar to the blue curve of HEXQ.

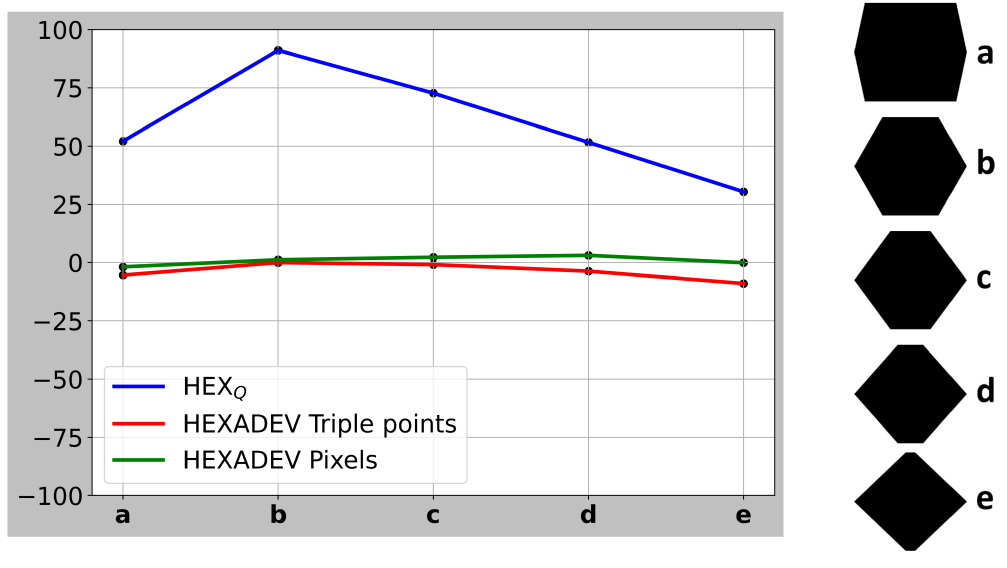

**Fig. S11.** Comparison of metrics assessing the similarity of a hexagon to a regular hexagon. Variation in the length of two opposing parallel sides.

Again, HEXQ exhibits more significant variations between different morphologies compared to the other criteria. Therefore, we observe that HEXADEV can present values in the range *[−100,+∞],* with practical values most often falling within the range *[−100, 0]*, as seen in the previous graphs or in the article [20]. Knowing that HEXQ ranges from 0 to 100, and based on the previous graphs, we can infer that HEXQ is slightly more sensitive than HEXADEV. HEXQ seems to better differentiate the various morphologies compared to the regular hexagon, justifying its relevance and its introduction in our study.
